# Steroid-resistant nephrotic syndrome associated *MYO1E* mutations have differential effects on myosin 1e localization, dynamics, and activity

**DOI:** 10.1101/2021.11.11.468158

**Authors:** Pei-Ju Liu, Laura K. Gunther, Diana Perez, Jing Bi-Karchin, Christopher D. Pellenz, Sharon E. Chase, Maria F. Presti, Eric L. Plante, Claire E. Martin, Svjetlana Lovric, Christopher M. Yengo, Friedhelm Hildebrandt, Mira Krendel

## Abstract

Myo1e is a non-muscle motor protein enriched in the podocyte foot processes. Mutations in *MYO1E* are associated with steroid-resistant nephrotic syndrome (SRNS). Here, we set out to differentiate between the pathogenic and neutral *MYO1E* variants identified in SRNS patients by exome sequencing. Based on protein sequence conservation and structural predictions, two mutations in the motor domain, T119I and D388H, were selected for this study. EGFP-tagged Myo1e constructs were delivered into the Myo1e-KO podocytes via adenoviral infection to analyze Myo1e protein stability, Myo1e localization, and clathrin-dependent endocytosis, which is known to involve Myo1e activity. Furthermore, truncated Myo1e constructs were expressed using the baculoviral expression system and used to measure Myo1e ATPase and motor activity *in vitro*. Both mutants were expressed as full-length proteins in the Myo1e-KO podocytes. However, unlike wild-type (WT) Myo1e, the T119I variant was not enriched at the cell junctions or clathrin-coated vesicles (CCVs) in podocytes. In contrast, the D388H variant localization was similar to the WT. Surprisingly, the dissociation of the D388H variant from cell-cell junctions and CCVs was decreased, suggesting that this mutation also affects Myo1e activity. The ATPase activity and the ability to translocate actin filaments were drastically reduced for the D388H mutant, supporting the findings from cell-based experiments. The experimental pipeline developed in this study allowed us to determine that the T119I and D388H mutations appear to be pathogenic and gain additional knowledge in the Myo1e role in podocytes. This workflow can be applied to the future characterization of novel *MYO1E* variants associated with SRNS.

## Introduction

Pediatric patients suffering from steroid-resistant nephrotic syndrome (SRNS) often carry genetic mutations, which undermine the selective permeability of the glomerular filtration barrier and result in proteinuria (excessive excretion of proteins in urine). SRNS often progresses to end stage renal disease (ESRD) that requires kidney transplantation. Patients with familial SRNS are good candidates for kidney transplantation due to low risk of disease recurrence [1–4]. To avoid the side effects of unnecessary steroid treatment, primary SRNS patients may need to be identified early and prioritized on a transplantation waiting list instead of taking steroids [5, 6]. One of the genes linked to primary SRNS is *MYO1E* (encodes Myo1e), which was initially examined in the familial cases of SRNS and exhibits the autosomal recessive pattern of inheritance of disease-associated variants [7]. With the expansion of exome sequencing, novel candidate mutations in the known SRNS genes, including *MYO1E*, have been identified [8–10]. Since identification of new variants by exome sequencing is not always accompanied by the corresponding family history, and the precise mechanisms linking Myo1e mutations to glomerular filtration disruption are unknown, it is important to develop a workflow for the functional characterization of novel *MYO1E* mutations to determine whether they are pathogenic and to predict their effects on myosin functions in the kidney. In this paper we set out to characterize novel mutations in *MYO1E* to differentiate pathogenic vs. neutral *MYO1E* variants among those identified by exome sequencing or other genomic methods [7–9].

Myo1e is an actin-dependent ATPase [11, 12] and a member of myosin class 1 [13]. Class 1 myosins bind actin filaments via their motor domains and plasma membrane via their tail domains, thus serving as actin-membrane linkers. Furthermore, some class 1 myosins, including Myo1e, contain additional protein interaction motifs in their tails that allow them to recruit additional binding partners to the actin-membrane interface. Myo1e is highly expressed in podocytes, terminally differentiated epithelial cells covering basement membrane atop of glomerular endothelium [7, 14, 15], and is enriched in podocyte foot processes, elongated protrusions that are connected to each other by slit diaphragms (modified tight/adherens junctions). Both podocyte foot processes and slit diaphragms are important components of the glomerular filter. Knockout of Myo1e (Myo1e-KO), either germline (complete) or podocyte-specific, leads to proteinuria in mice with accompanying foot process effacement (flattening) and irregular thickening of the glomerular basement membrane [14–16]. Similar to Myo1e-KO mice, patients with *MYO1E* mutations exhibit proteinuria along with podocyte foot process effacement and thickened glomerular basement membrane [7].

We have shown that Myo1e localizes at the podocyte cell-cell junctions/slit diaphragms and interacts with ZO-1 via its SH3 domain [7, 14, 17]. Myo1e SH3 domain also binds endocytic proteins dynamin and synaptojanin [18], and Myo1e is recruited to clathrin-coated vesicles immediately prior to vesicle scission, together with dynamin and actin [19]. Expression of a dominant-negative Myo1e construct or knockdown of Myo1e results in reduced receptor-mediated endocytosis in HeLa cells [18, 20]. Since knockouts of endocytic proteins Myo1e, dynamin, synaptojanin, and endophilin in podocytes lead to severe proteinuria in mice, clathrin-dependent endocytosis appears to play an important role in normal glomerular filtration [14, 21]. In addition, Myo1e (and a closely related myosin, Myo1f, which is not expressed in podocytes) regulate actin dynamics in macrophages [22]. Taken together, these observations suggest that Myo1e may contribute to maintaining slit diaphragm integrity or selective filtration via its roles in endocytosis, actin dynamics, or cell-cell contact assembly, and that mutations that affect Myo1e localization to cell-cell junctions, clathrin-coated vesicles, or actin filaments in podocytes are likely to be pathogenic.

Several novel *MYO1E* variants associated with childhood nephrotic syndrome have recently been found in a Saudi Arabian cohort and a global cohort of families with SRNS by exome sequencing [8, 9]. To investigate whether the novel Myo1e variants containing single amino acid substitutions are pathogenic, and to determine how these mutations affect Myo1e activity, we characterized protein expression/stability and intracellular localization of two of the Myo1e variants, T119I and D388H, using adenoviral expression of mutant constructs in the Myo1e knockout podocyte cell line. We found that the two point mutations in the myosin motor domain altered Myo1e localization and dynamics in podocytes. Specifically, the Myo1e^T119I^ variant I did not colocalize with actin filaments while the Myo1e^D388H^ variant exhibited reduced dissociation from actin bundles at cellcell junctions. In addition, expression of these myosin variants affected the density and lifetimes of clathrin-coated vesicles (CCVs) in podocytes. To gain additional insight into the effects of these mutations on Myo1e activity, we performed *in vitro* characterization of myosin motor activity using ATPase kinetics and filament sliding assays, with the latter providing the first characterization of the Myo1e-driven motility *in vitro.* The D388H mutation in the Myo1e motor domain led to a dramatic reduction of myosin ATPase activity and loss of motility, while the T119I variant was too unstable to be used for these measurements. Overall, our work reveals the deleterious effects of two disease-associated *MYO1E* genetic variants, provides a standardized workflow for characterization of novel *MYO1E* variants, and sheds light on the possible mechanisms causing defective glomerular filtration in SRNS patients.

## Methods

### Podocyte culture

Conditionally immortalized wild-type podocytes were a generous gift of Dr. Peter Mundel [23, 24] while conditionally immortalized Myo1e-KO podocytes were derived from the Myo1e-KO mice as described [14, 17]. Podocytes were grown on collagen I (Corning #354236) coated culture dishes in RPMI-1640 with 10%FBS, 1% antibiotic-antimycotic, and 50 μg/ml interferon-γ (EMD Millipore # 407303) at 33°C with 5% CO_2_. For differentiation, podocytes were cultured at 37°C without interferon-γ [23, 24]. Podocytes were differentiated for 10 days prior to adenoviral transduction for protein analyses and live-cell imaging following previously described transduction procedures [25].

### Constructs

Point mutations were introduced into the wild-type human Myo1e in the pEGFP-C1 vector using QuikChange Lightning Site-Directed Mutagenesis Kit (Agilent Technologies #210519) [7, 18]. Mutant constructs were used for subcloning into the pAdEasy adenoviral vector and packaged into recombinant adenoviral particles as described [17, 25]. For the baculoviral expression, an EGFP-FLAG-AviTag-encoding linear DNA segment was prepared by gene synthesis (GENEWIZ) and inserted into the pFastBac vector using In-Fusion kit (TaKaRa # 638910). Inserts encoding wild-type or mutant Myo1e fragments (motor domain + IQ motif, a.a.1-724) were PCR amplified from the pEGFP-C1-Myo1e constructs and subcloned by In-Fusion into the pFastBac/EGFP-FLAG-Avi vector between EGFP and FLAG. Primers used for cloning are listed in the Supplementary table 1.

### Protein expression in podocytes and drug treatments

To measure protein expression, Myo1e-KO podocyte lysates were collected 24 hrs post-infection. Protein degradation rate was measured after 3-hour treatment with 20μg/ml of cycloheximide (Sigma-Aldrich #C7698) in the complete RPMI medium. Protein accumulation rate upon proteasome inhibition was measured after 3-hour treatment with 10μM of MG-132 (Cell Signaling Technology #2194) in the RPMI medium.

### Real-time PCR

Adenoviral infection (the amount of adenoviral DNA introduced into cells) and expression of EGFP-Myo1e RNA were detected by SYBR Green based real-time quantitative PCR. Primer sequences are listed in the Supplementary table 1. 24 hours post-infection, podocytes were washed three times with warm 1× PBS and the viral DNA was extracted using DNeasy Blood & Tissue kit (Qiagen #69504) [26]. RNA isolation was done using TRIzol method (Invitrogen #15596026) and the cDNA synthesis (reverse-transcription PCR) was accomplished by using iScript cDNA Synthesis Kit (BioRad #1708890). The realtime PCR was done using iTaq Universal SYBR Green Supermix (BioRad #172-5121) and performed on BioRad CFX 384 Real Time PCR System.

### Western blotting

Cells were harvested by scraping in CHAPS lysis buffer (20mM Tris-HCl pH7.5, 500mM NaCl, 0.5% w/v CHAPS) with protease inhibitors (Thermo Scientific #A32965). Cells were pipetted up and down 30 times and left on the ice for 30 minutes. Cell lysates were then boiled with Laemmli sample buffer, and separated on 10-20% gradient SDS-PAGE gel, followed by transfer to PVDF. Membranes were blocked in 5% milk in TBST for 1 hour at room temperature. The primary antibody, rabbit anti-GFP (ThermoFisher Scientific #A6455), was diluted (1:2000) in 5% milk in TBST and incubated with the membranes overnight at 4°C. The next day, the membrane was washed 3 times for 5 minutes in TBST. The secondary antibody, goat anti-rabbit conjugated with HRP (Jackson ImmunoResearch), was diluted (1:5000) in 5% milk in TBST and incubated with the membranes for 1 hour at room temperature. Chemiluminescence was detected using WesternBright Quantum (Advansta) and imaged on a BioRad ChemiDoc imaging system.

### Podocyte culture for live-cell imaging

5X10^3^ podocytes were plated onto collagen IV (Corning #356233) coated 35mm glass-bottom MatTek (MatTek #P35G-15-14-C) for differentiation for 10 days. At day 10, podocytes were infected with recombinant adenoviruses. After 24 hours of adenoviral transduction, podocytes were washed with warm RPMI media 3 times and observed using confocal or total internal reflection fluorescence (TIRF) microscopy.

### Confocal microscopy

For cell-cell junction and fluorescence recovery after photobleaching (FRAP) analysis, images were taken using a Perkin-Elmer UltraView VoX Spinning Disk Confocal system mounted on a Nikon Eclipse Ti-E microscope and equipped with a Hamamatsu C9100-50 EMCCD camera, a Nikon Apo TIRF 60X (1.49 N.A.) oil objective, and controlled by Volocity software. An environmental chamber was embedded to maintain cells at 37°C. During imaging, cells were maintained in RPMI supplemented with 10%FBS and 1% antibiotics-antimycotics.

### TIRF microscopy

For clathrin-coated vesicle analysis, the cells were imaged using a TIRF illumination setup (Nikon) mounted on a Nikon Eclipse TE-2000E inverted microscope equipped with a perfect focus system (Nikon), a Nikon Apo TIRF 100X (1.49 N.A.) oil objective and Prime 95B camera (Photometrics) and controlled by NIS-Elements software. An environmental enclosure was used to maintain cells at 37°C. During imaging, cells were maintained in RPMI supplemented with 10% FBS and 1% antibiotic-antimycotic solution or in imaging buffer (136 mM NaCl, 2.5 mM KCl, 2 mM CaCl_2_, 1.3 mM MgCl_2_, and 10 mM HEPES [pH 7.4]). Both 488 nm and 561 nm laser power were set at 60%. Images were acquired every 5 seconds for 6 minutes, with exposures of 200 ms or 800 ms for the 561 nm and 800 ms for the 488 nm wavelength.

### Fluorescence intensity and kymographs

Fluorescence intensity measurements for junctional enrichment (Fig. 3 and S3), FRAP (Fig. 3 and S4), ROI mean fluorescence intensity (Fig. S3B-C and S5B-C) and endocytic puncta intensity (Fig. S6) as well as endocytic vesicle kymographs (Fig. 4D) were collected using Fiji [17, 20, 27, 28].

### FRAP imaging and analysis

For photobleaching, a 488 nm argon laser with full power was used to bleach a 30×75 pixel ROI at the podocyte junctions. 5 frames of pre-bleaching images were collected at 1 frame/sec. Post-bleaching images were acquired using the 488 nm laser at 25% of laser power; the acquisition rate for the first 40 sec was set to 2 frames/sec and for the subsequent 120 sec to 1 frame every 2 sec. Fluorescence intensity of the bleached area was measured over time and normalized relative to the background and a control region of interest (to correct for acquisition bleaching) as well as the prebleaching images. Lastly, the fluorescence intensity of the ROI at the first time point after bleaching was set to 0. The best-fit curve for fluorescence recovery was obtained using the exponential one-phase association model in Prism 8. The following equation was used: *y*=*a*(1-*e*^−*bx*^), where *x* is time in seconds. The half time of recovery was determined using *b* from the previous equation, where *t*_1/2_ = ln 0.5/−*b*. Analysis was performed on 16-bit images [27].

### Endocytic vesicle detection, tracking, and analysis

Quantification of the clathrin-coated vesicle (CCVs) and Myo1e co-localization (Fig. 4A), density (Fig. 4B), peak intensity (Fig. 4C) and lifetimes (Fig. 4E-F) was performed using Imaris software (v. 9.7.1, Bitplane Inc.). CCVs were automatically detected using the spot module and particle tracking following previously described procedures with additional modifications [29]. TIRF microscopy images were first subjected to ROI segmentation (to obtain images with an area of about 900 μm^2^) and background subtraction. CCVs were then detected using the spot module with an estimated spot diameter of 0.5 μm. The detected vesicles were filtered with Quality (defined as the intensity at the center of the spot, Gaussian filtered by ¾ of the spot radius). The threshold values of Quality were set to minimize the noise (as determined by visual inspection). Brownian motion particle-tracking algorithm was applied to trace objects through sequential frames of the time-lapse movies. If the distance between the candidate spot position and the predicted future position exceeded the maximum distance of 0.35 μm, the connections between a spot and a future position match were rejected. To connect tracks that were segmented due to the object temporarily out-of-focus, the maximum permissible gap length was set to 1 frame. Track outputs were then edited to correct for tracking errors by visual inspection. Only tracks that appeared from the first to the last frame (stable CCVs) as well as appeared and disappeared (dynamic CCVs) during the image acquisition (72 or 73 frames, 6 minutes) were subjected to lifetime analysis [29, 30].

Following the ROI segmentation for CCV tracking, Myo1e puncta were detected using the surface module (without creating tracks for the Myo1e particles). The puncta were first background subtracted, then the surface grain size was set to 0.1 μm and the diameter of the largest sphere was set to 0.28 μm. In addition, split touching objects (region growing) was enabled and seed point diameter was set to 0.4 μm. The thresholds for background subtraction, quality and number of voxels were set based on the visual inspection to minimize noise.

To collect the CCV-Myo1e co-localization information, object-object statistics in the CCV module was applied and CCV tracks were temporarily unlinked. Next, the shortest distance of CCV spot to Myo1e surface was filtered by setting the threshold to the maximum of 0.25 μm. Percentages of CCVs containing Myo1e (CCV-Myo1e co-localization rate, Fig.4A) were counted in a selected frame (the frame that is equivalent to the 90 sec time point in a 6 min movie). Further, the selected (Myo1e-colocalized) CCVs were tagged, and the data for mean intensity (Fig.4C), track duration (Fig.4E&F), and shortest distance to the Myo1e surface (Fig.4G&H) were extracted from the statistics section. Additionally, Gaussian filter (threshold=0.111) and local background subtraction (threshold=0.5μm) were applied prior to collecting CCV peak intensity.

CCV intensity (Fig.4C), lifetimes (Fig.4E&F), and association duration (Fig.4G&H) data collected from Imaris was first processed manually for statistical analysis. To reduce mistakes during repeated manual data processing, the data were then rearranged using Python code (https://github.com/fomightez/pjl), relying mainly on the Pandas module; the scripts were generously implemented by Wayne A. Decatur and provided within active Jupyter-based sessions served via MyBinder.org. For data rearrangement for CCV intensity analysis (Fig. 4C), maximum intensity value in each track was extracted and rearranged in a new list, with each list corresponding to a different experimental group for statistical analysis. For CCV lifetimes (Fig. 4E&F), in each experimental group, the values of track duration were moved to a new list for statistical analysis. For the association duration (Fig. 4G), the total frames of a track and the frames with the CCV-Myo1e distance of ≤0.25 μm were counted. In the resulting list, if the total number of frames was ≤5 (abortive CCVs), the track was removed from further analysis. Lastly, to calculate the duration of the Myo1e-CCV contact, the number of frames where Myo1e was associated with the CCV was divided by the total number of frames. The values of each experimental group were moved to a list for statistical analysis. Furthermore, identification of the CCVs that were only associated with Myo1e for a single frame was performed using C# code (https://github.com/opidopi/DataFilter) that was kindly created by Sean M. Lantry.

### Expression and purification of Myo1e for motor function assays

Recombinant baculoviruses expressing Myo1e constructs containing the motor and IQ domain as well as the C-terminal GFP, Avi, and FLAG tags were produced as described previously [31–33]. WT and D388H Myo1e were co-expressed with calmodulin in SF9 cells and purified with anti-FLAG affinity chromatography [34]. The purified Myo1e constructs were examined by Coomassie Blue staining of SDS-PAGE gels and concentrations were determined by GFP absorbance (ε_488_ = 55,000) and Bradford assays using BSA as a standard. Actin was purified from acetone powder derived from rabbit skeletal muscle (Pell-freeze) using the method of Pardee and Spudich [35].

### Steady-state ATPase measurements

The actin-activated ATPase activity of purified Myo1e was examined using the NADH-coupled assay [36] in KMg50 buffer (10 mM Imidazole, pH 7.0, 50 mM KCl, 1 mM EGTA, 2 mM MgCl_2_, and 1 mM DTT) at 25 °C. The absorbance of NADH was examined over a 200 second period in an Applied Photophysics Stopped-Flow apparatus. A standard curve of known ADP concentrations was used to determine the absorbance units per concentration of ADP. The ATPase rates (*V*) were plotted as a function of actin concentration, and the data were fit to a Michaelis-Menten equation (*V* = *V*0+(*k*_cat_*[actin])/(*K*_ATPase_+[actin]), which allowed determination of the maximum rate of ATPase activity, reported as the addition of the ATPase in the absence of actin (v_0_) and the actin-activated ATPase (k_cat_) (*V*_Max_ = *V*_0_+*k*_cat_), and actin concentration at which ATPase activity is one-half maximal (*K*_ATPase_). To remove non-functional Myo1e motors, we first bound Myo1e to actin, pelleted the actomyosin using ultracentrifugation (10 minutes at 95,000 RPMs in a TLA.120.2 rotor at 4 °C), and subsequently released the Myo1e from actin with a second ultracentrifugation step in the presence of 2 mM ATP and additional salt (final 150 mM KCl). The Myo1e released from actin in the presence of ATP was used directly in the ATPase and motility assays.

### Single nucleotide turnover

We used mant-labeled ATP (*mantATP*) to perform single ATP turnover experiments with FLAG purified WT and D388H Myo1e in the absence of actin. A sub-stoichiometric concentration of *mant*ATP was mixed with Myo1e in the stopped-flow apparatus, and mant fluorescence was examined over a 500 second period. The mant fluorescence, which increases when bound to myosin, was excited at 290 nm, and the emission measured with a 395 nm long-pass filter. The fluorescence decays were fit to the exponential equation that best fit the data.

### *In vitro* motility

The in vitro actin-gliding assay was performed as previously described [37, 38]. Myo1e was adhered to 1% nitrocellulose coated coverslips that contained anti-GFP antibody (0.1 mg/ml), and the surface was blocked with BSA (2 mg/ml). Sheared unlabeled actin (2 μM) followed by 2 mM ATP were added to block the dead myosin heads on the surface. The activation buffer containing KMg50 buffer, 0.35% methylcellulose, 0.45 mM phosphoenolpyruvate, 45 units/ml pyruvate kinase, 0.1 mg/ml glucose oxidase, 5 mg/ml glucose, 0.018 mg/ml catalase, and 2 mM ATP was added right before video acquisition. Alexa 555 phalloidin-labeled actin was visualized with a Nikon TE2000 fluorescence microscope equipped with a 60X/1.4NA lens and a Coolsnap HQ2 cooled CCD camera controlled using Nikon Elements 3.0. Videos were collected for up to 10 min at a 10 s frame rate. The velocities were manually analyzed by tracking actin filaments using MTrackJ in ImageJ [39].

### Protein sequence alignment and structure modeling

Protein sequence alignments were performed using ClustalW alignment function in MacVector (v. 17.0.10). The sequence accession numbers are listed in Supplementary table 2. Protein structure modeling and rotamer predictions were performed using UCSF Chimera [40].

### Statistical analysis

For multiple comparisons of the WT and mutants, data were analyzed using a one-way ANOVA with Tukey’s post-hoc test, with statistical significance set at p-value<0.05. All the statistical analyses and graphing were performed using GraphPad Prism software.

## Results

### *MYO1E* variants found in nephrotic syndrome patients and their predicted effects on protein structure

Exome sequencing of SRNS candidate genes in patients exhibiting SRNS and/or diagnosed with FSGS identified a number of variants in the *MYO1E* gene [8, 9]. Among the patients with non-truncating Myo1e mutations, patients homozygous for *MYO1E*^T119I^ or *MYO1E*^D388H^ presented with FSGS on renal biopsy and were resistant to steroid treatment **(Supplementary table 3)** [8, 9]. T119 and D388 are highly conserved residues in the myosin motor domain **(Fig. 1A-C)**. To determine if the mutated residues affect the key elements that are involved in myosin activity, we selected a class 1 myosin for which crystal structure has been determined, namely, the short-tailed *Dictyostelium* myosin myo1E (PDB: 1LKX) [41]. The T108 residue in myo1E (equivalent to the T119 in human Myo1e, **Fig. 1B**) is located within the P-loop, which, along with the switch-1, constitutes the structural elements essential for nucleotide binding [42] and sensing whether the bound nucleotide is ATP or ADP [43]. The threonine residue binds the magnesium ion associated with the ATP (**Fig. 1D)**. Upon changing the threonine at position 108 to isoleucine to model the effects of the T118I mutation, we found that the isoleucine side chain clashed with the magnesium ion and with the switch-1 residue S158 **(Fig. 1E)**. The D386 residue (equivalent to D388 in human Myo1e, **Fig. 1C)** is in the switch-2 region, which connects with the relay helix to convey the conformational changes to the converter domain during the ATPase cycle [44, 45]. Replacing D386 with a histidine, we modeled a number of possible histidine side chain orientations using UCSF Chimera. In rotamer 1 (probability score 0.18), the histidine clashed with S111 on the *α*-helix that connects to the P-loop **(Fig. 1F upper panel)**. In rotamer 2 (probability score 0.13), this histidine clashed with the T108 on the P-loop, which is the above-mentioned residue binding to the magnesium ion **(Fig. 1F lower panel)**. In both cases, the D386H mutation may disrupt the P-loop interaction with Mg-ATP. Thus, the T119I and D388H mutations may affect the ATPase activity of Myo1e, however, their precise effects on Myo1e activity and stability cannot be determined solely based on the modeling data. Therefore, we set out to experimentally test the effects of these mutations using cell biological and biochemical approaches.

**Figure 1.**
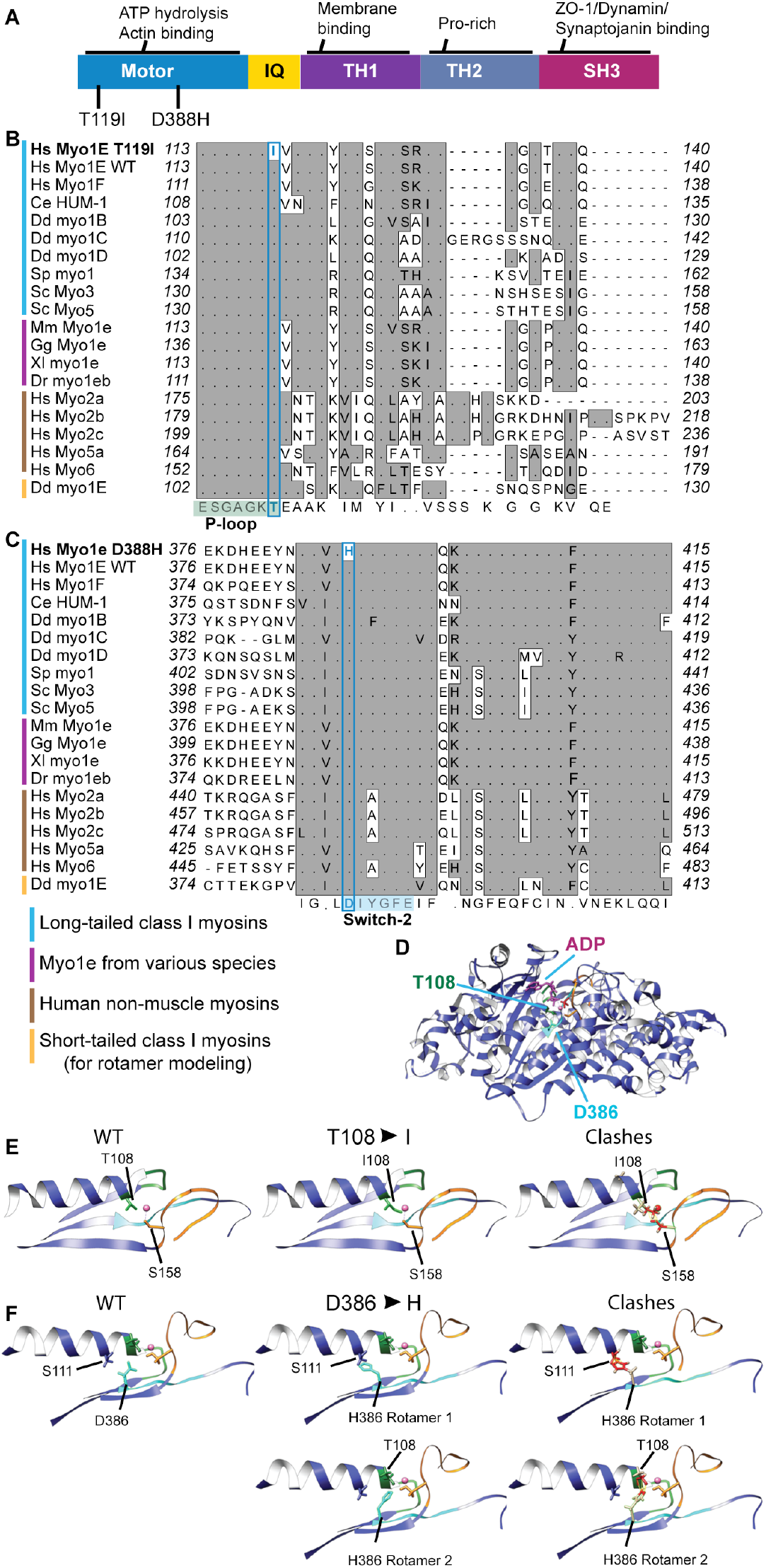
Amino acid residues T119 and D388 in the Myo1e motor domain are highly conserved. **(A)** Domain map of human Myo1e indicating the locations of the SRNS-associated Myo1e mutations T119I and D388H. (**B-C)** Alignment of myosin protein sequences for the regions containing T119 **(B)** and D388 **(C)** residues. Both amino acid residues are conserved among class 1 myosins, including long-tailed class 1 myosins from *Caenorhabditis elegans* (worm), *Dictyostelium discoideum* (slime mold), *Schizosaccharomyces pombe* (fission yeast), and *Saccharomyces cerevisiae* (budding yeast), the short-tailed class 1 myosin myo1E of *D. discoideum* that we used for the rotamer analysis, as well as in myosin 1e of *Mus musculus* (mouse), *Gallus gallus* (chicken), *Xenopus laevis* (frog) and *Danio rerio* (zebrafish). These residues are also conserved in other myosin classes, including human non-muscle myosin 2a-c, 5a, and 6. The conserved residues corresponding to the mutation site are highlighted in blue boxes. **(D)** Ribbon diagram representation of the ADP-bound *D. discoideum* myo1E motor domain (PDB: 1LKX) showing the residues T108 and D386 that are equivalent to the human Myo1e residues T119 and D388. Their locations are shown relative to the key structural elements of the magnesium-binding site. The backbone of the myo1E structure is shown as a blue ribbon. P-loop is highlighted as a green ribbon. Switch-1 is highlighted as an orange ribbon. Switch-2 is highlighted as a cyan ribbon. Magnesium is highlighted in pink. *D. discoideum* myo1E residue numbers are shown. **(E)** A more detailed view of the T108 residue (green). Predicted structure of the most likely rotamers of I108 shows that one of the likely rotamers clashes with a switch I residue, S158. **(F)** A more detailed view of the D386 residue (cyan). Predicted structure of the most likely rotamer of the H386 shows that one of the likely rotamers clashes with S111 on an alpha helix connecting with the P-loop **(F, upper panel)**. Predicted structure of the second likely rotamer of the H386 shows that one of the likely rotamers clashes with T108 on the P-loop **(F, lower panel)**.

### Protein stability of the SRNS-associated Myo1e variants

To investigate whether the T119I and D388H mutations affect Myo1e protein stability, we introduced N-terminally-EGFP-tagged human Myo1e constructs into immortalized Myo1e-null mouse podocytes (KO podocytes) using adenoviral vectors and verified successful viral delivery of the Myo1e constructs into podocytes using isolation of viral DNA followed by qPCR **(Fig. S1A)**. Blotting with the anti-GFP antibody revealed the presence of the full-length proteins corresponding to the EGFP-myo1e fusions in the Myo1e^WT^-, Myo1e^T119I^-, and Myo1e^D388H^-expressing cells, and there was no significant decrease in the protein expression level among mutant constructs **(Fig. 2A&B, S1B)**. Additionally, using qRT-PCR, we found that the RNA expression for the mutants was 50 times higher than for the Myo1e^WT^ using *GAPDH, HPRT1* and *RPLP0* as housekeeping gene controls **(Fig. S1C)**. This indicates that the increased mRNA expression/stability could potentially compensate for any changes in protein stability but overall, no decrease in mutant protein expression was detected. To test whether there was a difference in the rate of protein turnover of mutant vs. wild-type proteins, we treated the Myo1e expressing podocytes with cycloheximide (CHX), a ribosome inhibitor, to stop protein expression and measure Myo1e degradation over time. Mutant Myo1e protein degradation rates were not significantly different from the Myo1e^WT^**(Fig. 2C&D, S1D)**. In addition, we inhibited proteasomal protein degradation using MG-132 and measured the accumulation rate of Myo1e^WT^ and variants. Protein accumulation rates of Myo1e^T119I^ or Myo1e^D388H^ were not significantly different from the Myo1e^WT^ when expressed in the Myo1e-KO podocytes **(Fig. 2E&F, S1E)**. On the other hand, when protein expression level and protein turnover were examined using expression of EGFP-tagged Myo1e in WT mouse podocytes or in HEK-293 human cells, Myo1e^T119I^ was found to be expressed at a lower level than Myo1e^WT^ and was rapidly degraded **(Fig. S2).** Similarly, our attempts to express and purify Myo1e^T119I^ in the baculovirus system resulted in very low protein yield and significant protein degradation, indicating a decreased stability of this protein. Myo1e^D388H^ protein was less stable than Myo1e^WT^ when expressed in the baculovirus system but did not exhibit increased degradation in the wild-type podocytes or HEK-293 cells. Overall, while the SRNS-associated Myo1e^T119I^ and Myo1e^D388H^ variants appear to be expressed as full-length proteins in KO podocytes, with no evidence of increased degradation, these mutations may affect Myo1e protein stability in other cell types or in the presence of the wildtype Myo1e protein.

**Figure 2.**
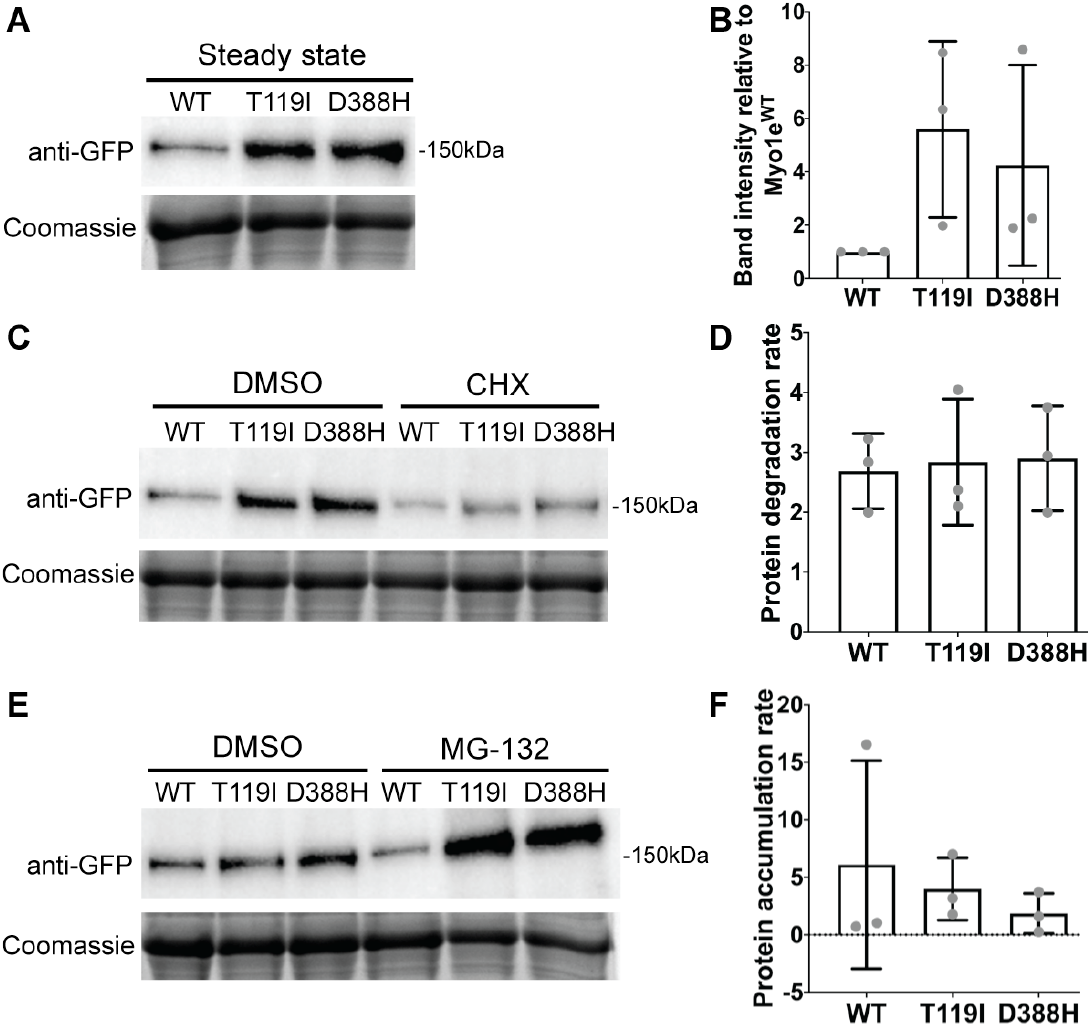
Myo1e^T119I^ and Myo1e^D388H^ are expressed as full-length proteins in Myo1e-KO podocytes. **(A)** Western blot analysis of total cell lysates of Myo1e-KO podocytes expressing Myo1e^WT^, Myo1e^T119I^ and Myo1e^D388H^ at steady state. (**B)** Quantification of band intensity in (A), normalized to the WT (mean±SD). (**C)** Western blot analysis of total cell lysates of Myo1e-KO podocytes expressing Myo1e^WT^, Myo1e^T119I^ and Myo1e^D388H^ treated with 20 μg/ml cycloheximide (CHX) or DMSO for 3 hours. (**D)** Myo1e degradation rate (band intensity in the DMSO-treated sample divided by the band intensity in the CHX-treated sample for the same construct) after CHX treatment in KO podocytes (mean±SD). (**E)** Western blot analysis of total cell lysates of Myo1e-KO podocytes expressing Myo1e^WT^, Myo1e^T119I^ and Myo1e^D388H^ treated with DMSO or 10 μM MG-132 for 3 hours. (**F)** Myo1e accumulation rate (band intensity in the MG-132-treated lysates divided by band intensity in the DMSO control) after the MG-132 treatment (mean±SD). **(A, C, E)** Blots were probed with the anti-GFP antibody. Equal protein loading verified by Coomassie Blue staining. **(B, D, F)** Data collected from 3 independent experiments. There is no significant difference (P>0.05) among groups as determined by one-way ANOVA.

### Myo1e^T119I^ localization to podocyte cell-cell junctions is disrupted

We have previously found that Myo1e localized to cellcell junctions in podocytes and played a role in regulating junctional assembly [7, 17]. Therefore, we set out to determine whether the recently identified SRNS-associated mutations affect Myo1e localization to cell-cell contacts in Myo1e-KO podocytes. EGFP-Myo1e^WT^ co-localized with the junctional marker mCherry-ZO-1 **(Fig. 3A)**. Myo1e^T119I^ was mostly absent from cell-cell contacts but such conspicuous mis-localization was not observed for the Myo1e^D388H^ **(Fig. 3A)**. We quantified junctional Myo1e enrichment **(Fig. 3B, S3A)**. This quantitative analysis confirmed the lack of localization of Myo1e^T119I^ at the cell-cell contacts while junctional localization of Myo1e^D388H^ in KO podocytes was not affected **(Fig. 3B)**. To find out whether the varying levels of EGFP-Myo1e and ZO-1-mCherry expression between cells could affect junctional enrichment measurements, we checked whether Myo1e junctional enrichment correlated with the level of Myo1e or ZO-1 expression in each cell. Using linear regression analysis, we found no significant positive correlation between the mean fluorescence intensity (MFI) of ZO-1-mCherry or EGFP-Myo1e and the extent of junctional enrichment in each Myo1e variant **(Fig. S3B&C),** indicating that differences in expression level between individual cells likely do not affect the outcome of this analysis.

**Figure 3.**
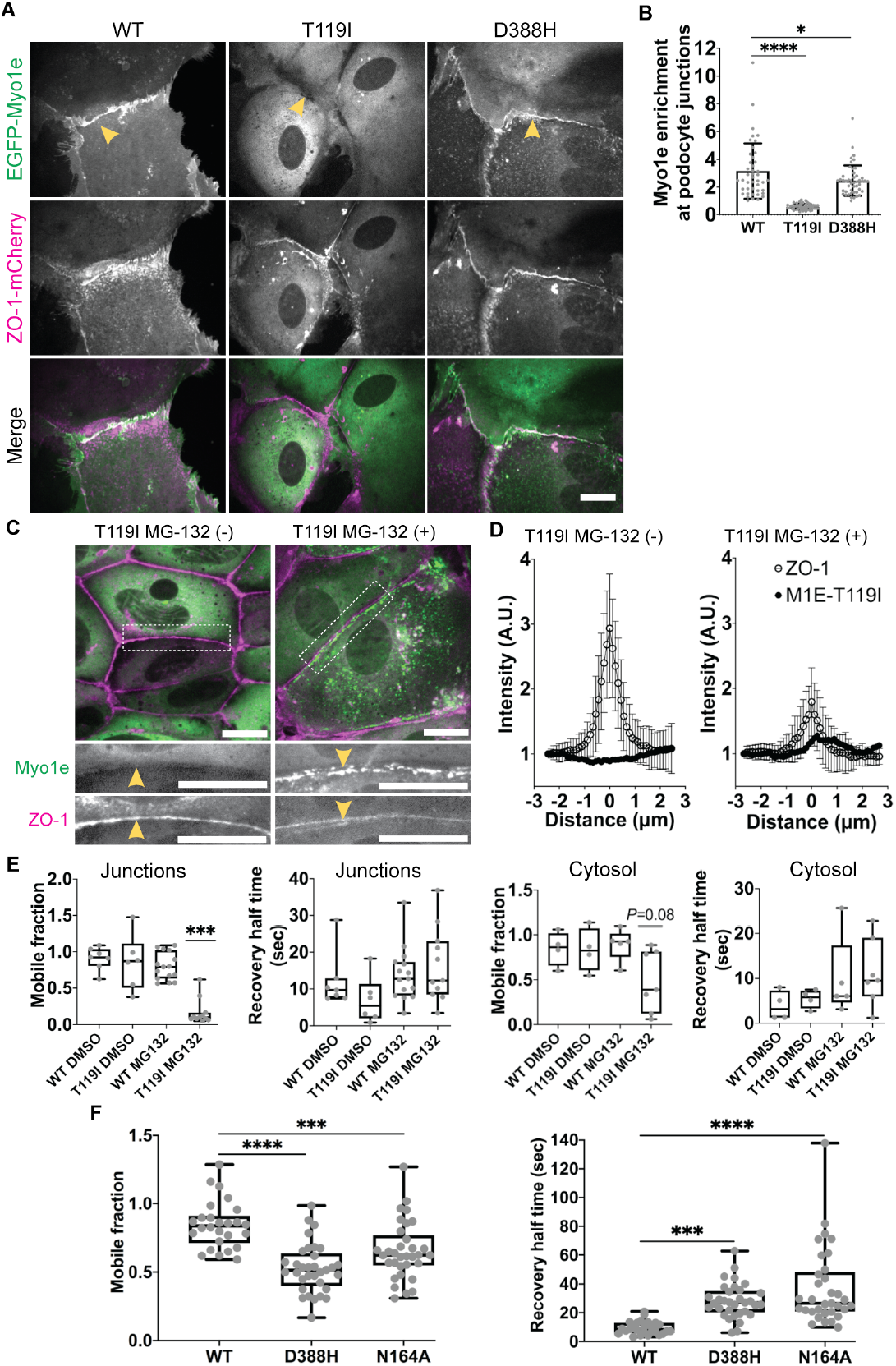
Junctional localization of Myo1e^T119I^ is disrupted and junctional protein exchange of Myo1e^D388H^ is decreased in podocytes. **(A)** Single confocal sections of Myo1e-KO podocytes co-expressing ZO-1-mCherry and EGFP-Myo1e^WT^, -Myo1e^T119I^ or -Myo1e^D388H^. Yellow arrowheads indicate cell-cell junctions. Scale bar, 20 μm. **(B)** Bar graph of Myo1e enrichment at the cell-cell junctions (mean±SD). 44 Myo1e^WT^-, 51 Myo1e^T119I^ - and 47 Myo1e^D388H^-expressing cells from 2 to 3 independent experiments were quantified. **(C)** Merged single confocal sections of KO podocytes co-expressing EGFP-Myo1e^T119I^ and ZO-1-mCherry after 10 μM MG-132 treatment for 3 hours. Dotted white boxes indicate the zoomed-in view shown in the lower panels. Yellow arrowheads point to podocyte junctions. Scale bar, 20 μm. **(D)** Quantification of junctional protein fluorescence intensity in **(C)** by line scan. Line scans of junctions without **(D, left)** and with **(D, right)** MG-132 treatment. Hollow circles indicate mean fluorescence intensity of ZO-1-mCherry and solid circles indicate mean fluorescence intensity of EFGP-Myo1e^T119I^ (mean±SD). 40 MG-132 (-) cells and 26 MG-132 (+) cells from 1 independent experiment were quantified. **(E)** Graphs of mobile fraction and recovery half-time of EGFP-Myo1e^T119I^ at cell-cell junctions **(E, 2 left panels)** and in the cytosol **(E, 2 right panels)** with DMSO or 10 μM MG-132 treatment for 3 hours. DMSO treated junctional FRAP was quantified from 7 Myo1e^WT^- and 6 Myo1e^T119I^-expressing cells. MG-132 treated junctional FRAP was quantified from 15 Myo1e^WT^- and 11 Myo1e^T119I^-expressing cells. DMSO treated cytosolic FRAP was quantified from 4 Myo1e^WT^- and 4 Myo1e^T119I^-expressing cells. MG-132 treated cytosolic FRAP was quantified from 5 Myo1e^WT^- and 7 Myo1e^T119I^-expressing cells. Data from 1 to 2 independent experiments. **(F)** Graphs of mobile fraction and recovery half-time of EGFP-Myo1e^WT^, Myo1e^D388H^, and Myo1e^N164A^ at cell-cell junctions at steady state. 26 Myo1e^WT^-, 33 Myo1e^D388H^- and 33 Myo1e^N164A^-expressing cells from 2 to 3 independent experiments were quantified. **(E-F)** Box and whisker plots indicate the median value, interquartile range, and full range of data points. Asterisks indicate significant difference from the Myo1e^WT^ control as determined by one-way ANOVA, ***P≤0.001 and ****P<0.0001.

Since Myo1e^T119I^ protein level was elevated after proteasomal inhibition **(Fig. S1E)**, we tested if increasing mutant protein availability would enhance its localization to the cell-cell junctions. After the MG-132 treatment, Myo1e^T119I^ (but not Myo1e^WT^) formed patchy clumps in the cytosol that, in some cases, aligned with cell-cell contacts **(Fig. 3C, S3D)**. Using line scans to compare the precise positions of the fluorescence intensity peaks of EGFP-Myo1e^T119I^ and ZO-1-mCherry, we found that the EGFP-Myo1e^T119I^ that accumulated upon proteasomal inhibition was not co-localized with ZO-1 **(Fig. 3D)**. We hypothesized that Myo1e^T119I^ patches may represent misfolded/immobilized protein aggregates rather than Myo1e complexes with ZO-1 that normally form at cell-cell junctions. To test this hypothesis, we measured protein dynamics of EGFP-Myo1e^T119I^ at the cell-cell junctions or in the cytosol of KO podocytes after the MG-132 treatment using fluorescence recovery after photobleaching (FRAP). Compared to the Myo1e^WT^ or DMSO-treated Myo1e^T119I^ cells, Myo1e^T119I^ puncta at the junctions had lower mobile fraction and slower fluorescence intensity recovery rate after the MG-132 treatment **(Fig. 3E)**. Likewise, the mobile fraction in the cytosol of Myo1e^T119I^-expressing cells was decreased after the MG-132 treatment **(Fig. 3E)**. In addition, we separated the MG-132 treated cell lysates into the supernatant and pellet fractions and found that Myo1e^T119I^ and Myo1e^D388H^ had lower solubility than Myo1e^WT^, and the majority of the protein remained in the pellet fraction **(Fig. S3E&F)**. Thus, when the misfolded Myo1e^T119I^ protein is prevented from being degraded by proteasomes, it forms immobile protein aggregates, which often accumulate near cellcell junctions.

### Myo1e^D388H^ localizes to cell-cell junctions but exhibits decreased dissociation from the junctions

Myo1e^D388H^ localized to cell-cell junctions, with a similar localization pattern to Myo1e^WT^ **(Fig. 3A&B)**. We examined the dynamics of Myo1e^D388H^ dissociation from cell-cell contacts using FRAP analysis **(Fig. 3F, Fig. S4)** and used an engineered mutant, Myo1e^N164A^, which was designed to be a rigor mutant with strong actin binding [46], as a positive control for FRAP experiments. Both Myo1e^D388H^ and Myo1e^N164A^ localized to cellcell junctions **(Fig. S4A)** and exhibited decreased mobile fraction and extended recovery half-time **(Fig. 3F, S4B&C, Supplementary movie 1**). Thus, Myo1e^D388H^ point mutation in the motor domain changes Myo1e-actin binding or other proteinprotein interactions at cell-cell junctions, resulting in slower protein exchange.

### Expression of Myo1e^T119I^ or Myo1e^D388H^ affects CCV dynamics

To test whether Myo1e localization to CCVs in podocytes is affected by the SRNS-associated mutations, we co-expressed mCherry-tagged clathrin light chain (CLC-mCherry) and EGFP-tagged Myo1e constructs in KO podocytes and used total internal reflection fluorescence microscopy (TIRFM) to image CCVs at the plasma membrane **(Fig. S5A)**. First, we examined the extent of co-localization of Myo1e with CLC by determining the % of all CLC-positive vesicles that were also Myo1e positive at the selected time point in cells expressing Myo1e variants. Consistent with the analysis of junctional enrichment, the extent of CCV co-localization with Myo1e was significantly lower for the Myo1e^T119I^ than for the Myo1e^WT^ and Myo1e^D388H^ **(Fig. 4A)**. There was no significant positive correlation between the MFI of CLC-mCherry or EGFP-Myo1e and the extent of co-localization in each Myo1e variant **(Fig. S5B&C),** indicating that differences in expression level between individual cells likely do not affect the outcome of the colocalization analysis.

**Figure 4.**
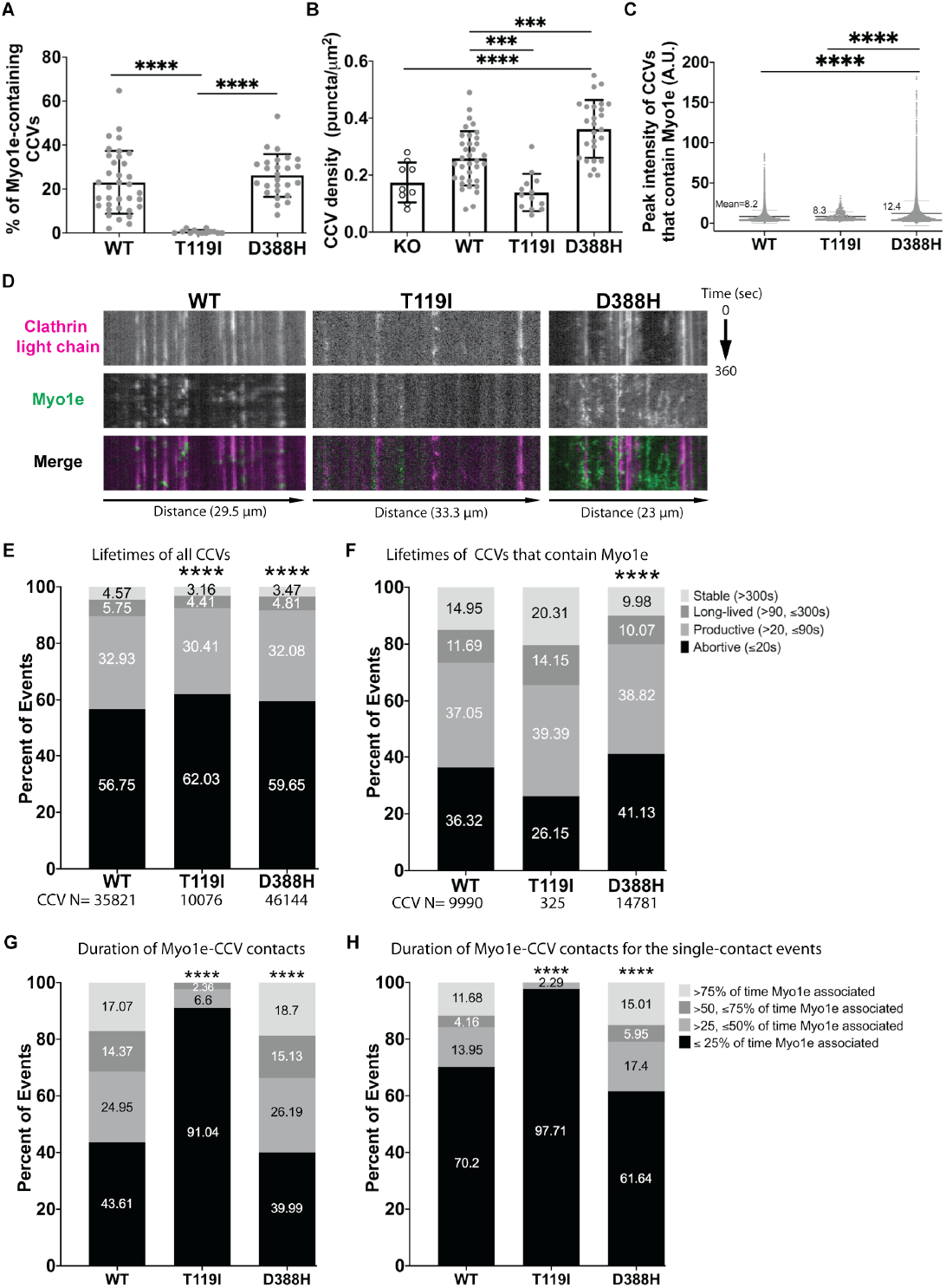
Myo1e^T119I^ and Myo1e^D388H^ expression in place of Myo1e^WT^ changes clathrin-coated vesicle (CCV) density, peak intensity, and lifetimes in podocytes. **(A)** Percent (mean±SD) of CCVs that contain Myo1e puncta in KO podocytes in a single time point image. **(B)** Density (mean±SD) of CCV (puncta/μm^2^) in KO podocytes expressing Myo1e mutant constructs. **(A-B)** 9 KO (CLC alone, no Myo1e expression), 35 Myo1e^WT^-, 13 Myo1e^T119I^- and 26 Myo1e^D388H^-expressing cells from 1-3 independent experiments were quantified. **(C)** Peak fluorescence intensity (mean±SD) of the CCV tracks that contain Myo1e measured after local background subtraction. Mean value of each experimental group is shown. 28 Myo1e^WT^-, 13 Myo1e^T119I^- and 26 Myo1e^D388H^- expressing cells from 2-3 independent experiments were quantified. **(D)** Kymographs showing CCV density, dynamics and Myo1e recruitment in Myo1e^WT^-, Myo1e^T119I^-, and Myo1e^D388H^-expressing KO podocytes. **(E)** Distribution of CCV lifetimes. **(F)** Distribution of the lifetimes of CCVs that contain Myo1e. **(E-F)** Lifetimes of CCVs were measured and categorized into abortive (≤20 sec, shown as 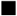), productive (>20 sec but ≤90 sec, shown as 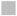), long-lived (>90 sec but ≤300 sec, shown as 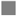), stable (>300 sec, shown as 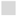). **(G)** Duration of Myo1e association with CCVs expressed as a fraction of the CCV lifetime. Only subpopulations with the lifetimes >20s were analyzed. **(H)** Duration of Myo1e association with CCVs for the CCVs that exhibited only a single Myo1e association event and CCV lifetimes >20s. **(E-F)** Duration of Myo1e association was measured and categorized into quartiles (≤25% of CCV track duration, shown as 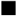; >25 but ≤50% of CCV track duration, shown as 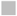; >50 but ≤75% of CCV track duration, shown as 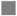; >75% of CCV track duration, shown as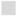). **(E-H)** Percent values of events are indicated on the bar plots. 28 Myo1e^WT^-, 13 Myo1e^T119I^- and 26 Myo1e^D388H^-expressing cells from 2 to 3 independent experiments were quantified. Asterisks indicate a significant difference from Myo1e^WT^ as determined by one-way ANOVA, 8**P≤0.001 and ****P≤0.0001.

Interestingly, the density of CLC puncta (number of vesicles per unit area) was significantly reduced in the KO podocytes expressing Myo1e^T119I^ and significantly increased in cells expressing Myo1e^D388H^ compared to the KO podocytes expressing Myo1e^WT^ **(Fig. 4B)**. The CCV density in the KO podocytes expressing Myo1e^T119I^ was similar to the KO podocytes not infected with Myo1e (Myo1e-null) **(Fig. 4B)**. To determine whether Myo1e affects the recruitment of clathrin to the endocytic pits, CLC intensity in the CCVs that contained Myo1e was measured over time. Peak clathrin intensity was D388H significantly increased in the Myo1e^D388H^-expressing KO podocytes, but no change was found in Myo1e^T119I^-expressing cells compared to the Myo1e^WT^-expressing cells **(Fig. 4C, S6F)**. In addition, five Myo1e-containing CLC puncta in each cell were manually selected, and their fluorescence intensity was measured using a line scan **(Fig. S6A**) to obtain both intensity plots **(Fig. S6C-E)** and intensity values at the center of each structure **(Fig. S6B)**. Consistent with the peak CLC intensity data, the central intensity of CLC was significantly increased in the Myo1e^D388H^-expressing KO podocytes **(Fig. S6B-E)**, suggesting that mutant Myo1e may affect the recruitment of clathrin or stabilization of the CCVs at the membrane. Together with the observation that CCV density is affected by the Myo1e mutant expression, our findings suggest that Myo1e not only influences vesicle internalization or scission, as previously suggested, but may also affect CCV stabilization during the earlier steps in endocytosis, affecting the number and size of CCVs.

Myo1e is recruited to endocytic invaginations immediately before the internalization of CCVs [19]. While Myo1e^WT^ was transiently recruited to CCVs immediately before their disappearance from the TIRFM image, Myo1e^T119I^ localization was diffuse, and very few instances of its transient recruitment to CCVs were observed **(Fig. 4D)**. On the other hand, Myo1e^D388H^ formed longer-lived and numerous puncta, some of which were co-localized with CCVs **(Fig. 4D)**. To perform a functional test of clathrin-dependent endocytosis, we measured uptake of transferrin, a prototypical clathrin-dependent endocytic cargo, in podocytes. However, mouse podocytes did not take up either human or mouse transferrin. In addition, to test whether Myo1e affects actin recruitment to the CCVs, we used Lifeact or siR-actin to image F-actin at the CCVs in live cells. However, we found that CCV-associated actin was not efficiently labeled by the reagents tested, likely due to low efficiency of Lifeact and siR-actin binding to the rapidly assembling, transient actin structures [47, 48].

To determine whether the expression of mutant Myo1e affects CCV assembly, maturation, and/or departure from the membrane, we tracked CLC and Myo1e in time-lapse series [20, 49]. Tracking CLC-mCherry, we classified CCV lifetimes into abortive (≤20 sec), productive (>20 sec but ≤90 sec), long-lived (>90 sec but ≤300 sec) and stable (>300 sec) [20, 49]. Both Myo1e^T119I^ - and Myo1e^D388H^-expressing cells contained higher fraction of abortive CCVs than Myo1e^WT^-expressing cells **(Fig. 4E).** We then examined separately the lifetimes of CCVs that contained Myo1e; they were characterized by a lower proportion of abortive CCVs than the general CCV population **(Fig. 4F).** There was a further decrease in the abortive CCV population and an increase in the stable CCV population among Myo1e^T119I^ - positive CCVs compared to the Myo1e^WT^-positive CCVs **(Fig. 4F)**, although this analysis may be complicated by the low number of the Myo1e^T119I^-positive CCVs **(Fig. 4A)**. On the other hand, in the Myo1e^D388H^-expressing cells, the proportion of abortive CCVs was elevated and that of the stable population was reduced **(Fig. 4F)**.

We next measured the duration of Myo1e association with the CCVs as a fraction of each CCV’s lifetime and categorized the resulting values into quartiles. Intriguingly, in the Myo1e^T119I^ expressing cells, 91% of the Myo1e-containing CCV were associated with Myo1e for ≤25% of the track lifetime, while this short-duration fraction constituted only 43.61% of the CCVs in the Myo1e^WT^-expressing cells **(Fig. 4G)**. Furthermore, in some cases Myo1e was recruited to a single CCV track multiple times. Because the nature of these repeated Myo1e-CCV interactions is unclear, we excluded these CCVs from our analysis and reanalyzed Myo1e association for those CCVs that exhibited only a single event of Myo1e recruitment **(Fig. 4H)**. With this analysis, the Myo1e^D388H^ was characterized by a higher proportion of the CCVs that exhibited prolonged association with Myo1e compared to the Myo1e^WT^ **(Fig. 4H)**. Overall, our analysis indicates that Myo1e^T119I^ is not recruited to CCVs to the same extent as the Myo1e^WT^, and that expression of this variant affects CCV density and internalization while Myo1e^D388H^ is characterized by the prolonged retention at the sites of clathrin-dependent endocytosis.

### SRNS-associated mutations affect Myo1e motor activity

We then set out to purify myosin constructs to measure ATPase and motor activity *in vitro.* To obtain sufficient amounts of Myo1e for these assays, we turned to the baculovirus expression system, which has previously been used to express a truncated Myo1e protein consisting of the motor and neck domain with the addition of a C-terminal FLAG tag for affinity purification [50]. In this earlier study, the truncated Myo1e protein was used for the kinetic characterization of the actin-activated ATPase activity. We added two additional C-terminal tags for motility assays: an EGFP tag that could elevate the myosin motor domain above the coverslip to avoid steric interference with the motor activity and an Avi-tag that could be used to couple the motor to the coverslip via biotin-streptavidin bonds **(Fig. 5A)**. Using the Myo1e^WT^ construct, we found that attachment of the EGFP-tagged protein to the coverslip using an anti-GFP antibody supported robust motor activity *in vitro*, and that biotinylation was not needed to provide strong adhesion to the motility chamber.

**Figure 5.**
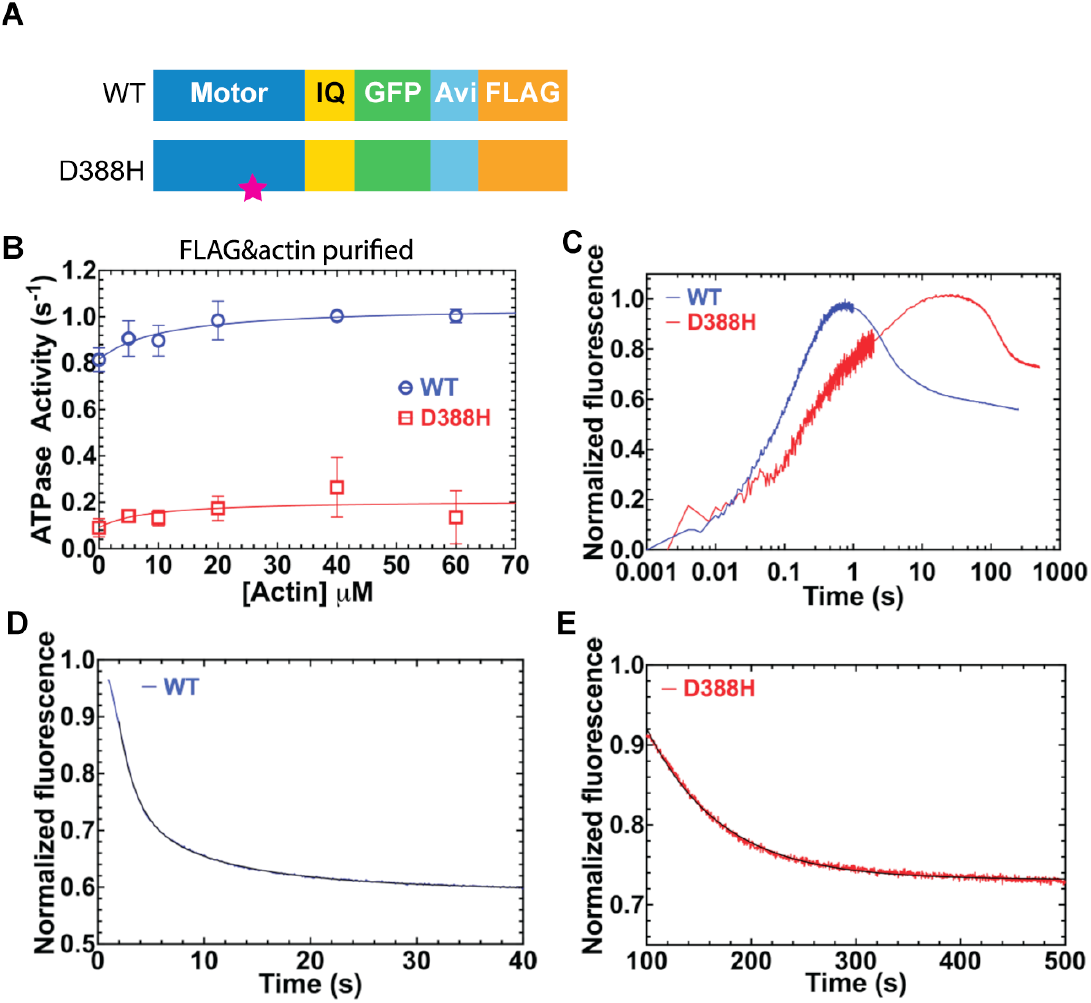
D388H mutation in Myo1e disrupts motor ATPase activity. **(A)** Myo1e construct design for baculovirus expression. **(B)** Actin-dependent ATPase activity of the FLAG- and actin-co-sedimentation-purified Myo1e constructs was measured using the NADH-coupled ATPase assay. The data points represent the mean±SD from 3 independent experiments for both WT and D388H. The solid lines are best fit to a Michaelis-Menten model. **(C)** Single ATP turnover kinetics using FLAG purified Myo1e in the absence of actin. Sub-stoichiometric amount of *mantATP* was mixed with Myo1e. We observed a fluorescence increase that was followed by an exponential decrease, which was used to estimate the single turnover rate constant. **(D)** The fluorescence decay of Myo1e^WT^ was fit to a three-exponential equation, **(E)** while the decay of Myo1e^D388H^ was single exponential (Table 2). The single turnover data were collected from a single experiment for both WT and D388H.

Due to the susceptibility of the Myo1e^T119I^ to proteolysis, we were not able to purify enough of this protein to perform *in vitro* assays. Myo1e^D388H^ was more stable but still exhibited elevated degradation compared to the Myo1e^WT^ **(Fig. S7A)**. To preclude the interference of degraded protein in the assessment of motor kinetics, the proteins were subjected to additional enrichment for active motors using actin binding followed by ATP release. The proteolytic bands remained even after the additional actin-binding purification in both Myo1e^WT^ and Myo1e^D388H^, albeit the latter showed more proteolysis than the former **(Fig. S7A)**, therefore, we used both actin-purified and standard myosin preparations in our experiments.

To characterize the kinetics of Myo1e, we measured its ATPase activity in the presence of varying actin concentrations **(Fig. 5B, Fig. S7B)**. In the absence of actin, Myo1e^WT^ exhibited relatively high basal ATPase activity (*V*_0_=0.82±0.05 s^−1^), which was observed in a previous study [50] and is unusual for myosins. In the presence of actin the ATPase was maximally increased approximately 30% (VMAX = 1.05±0.08 s^−1^). The basal ATPase activity of Myo1e^D388H^ was substantially reduced (*V*0=0.09±0.04 s^−1^) compared to Myo1e^WT^. However, the ATPase activity was activated approximately 2-fold in the presence of actin (V_MAX_ = 0.21±0.07). Overall, the maximum ATPase rate (*V*_MAX_) of Myo1e^WT^ was 5 times higher than that of Myo1e^D388H^ The actin concentration needed to achieve half-maximum ATPase activity (*K*_ATPase_ (μM)) was similar in WT and mutant Myo1e **(Fig. 5B, Table 1)**. The measurements of Myo1e ATPase activity were also examined without the extra step of purifying by actin co-sedimentation (FLAG purified), which demonstrated that the actin co-sedimentation step slightly increased the maximum ATPase activity but did not change the relative differences observed between Myo1e^WT^ and Myo1e^D388H^ **(Fig. S7B&C)**. To further examine the ATPase kinetics in the absence of actin, single ATP turnover experiments were performed on FLAG purified Myo1e. By mixing Myo1e with fluorescently-labeled ATP (*mant*ATP) we observed a fluorescence increase, associated with *mant*ATP binding to myosin, followed by a fluorescence decay, which should represent the single turnover rate constant. We found that the fluorescence decay was best fit by a single exponential for Myo1e^D388H^ but was triphasic for Myo1e^WT^ **(Fig. 5C-E)**. The results showed that the fast phase of the fluorescence decay in Myo1e^WT^ is most similar to the basal ATPase rate constant while the slower phases may represent a different conformation of Myo1e that releases products slowly. Myo1e^D388H^ was able to bind and hydrolyze ATP and the single turnover rate constant was relatively similar to the basal ATPase rate measured in the steady-state ATPase assay **(Fig. 5C-E, Table 2)**.

**Table 1.** Measurements of the ATPase activity of the Myo1e motor-IQ constructs (actin co-sedimentation purified).

| Construct | $V_0$ (s<sup>-1</sup>) | $V_{MAX}$ (s<sup>-1</sup>) | $K_{ATPase}$ (μM) |
| --- | --- | --- | --- |
| WT (N=3) | 0.82±0.05 | 1.05±0.08 | 10.7±7.5 |
| D388H (N=3) | 0.09±0.04 | 0.21±0.07 | 8.2±14.3 |

**Table 2.** Measurements of the single turnover kinetics of the Myo1e motor-IQ constructs.

| Construct | $k_{Fast}$ (s<sup>-1</sup>) | $k_{Slow}$ (s<sup>-1</sup>) |
| --- | --- | --- |
| WT (N=1) | 0.52±0.01 | 0.14±0.01 & 0.06±0.01 |
| D388H (N=1) | NA | 0.014±0.001 |

To investigate the functional consequences of the reduced ATPase activity in Myo1e^D388H^, we performed *in vitro* motility measurements with the Myo1e motor-IQ constructs (WT and D388H) **(Fig. 6)**. Using comparable motor densities, Myo1e^WT^ supported robust F-actin sliding (V_avg_=117.9±49.4 nm/sec, at 1 μM Myo1e surface density) **(Fig. 6A, Table 3, Supplementary movie 2)**, whereas, no F-actin translocation was detected in the assays with the Myo1e^D388H^ preparations at any density **(Table 3, Supplementary movie 2)**. When analyzing the motor density dependence of the Myo1e^WT^-mediated motility, we observed some variability but overall similar velocities at varying motor densities **(Fig. 6B).** Finally, mixing 1μM Myo1e^WT^ with varying amounts of Myo1e^D388H^, we found that the observed velocity of F-actin decreased with the increasing amount of Myo1e^**D388H**^. This indicates that Myo1e^D388H^ can bind to F-actin and slow Myo1e^WT^ motility, essentially functioning as a motor dead mutant **(Fig. 6C, Supplementary movie 2)**.

**Figure 6.**
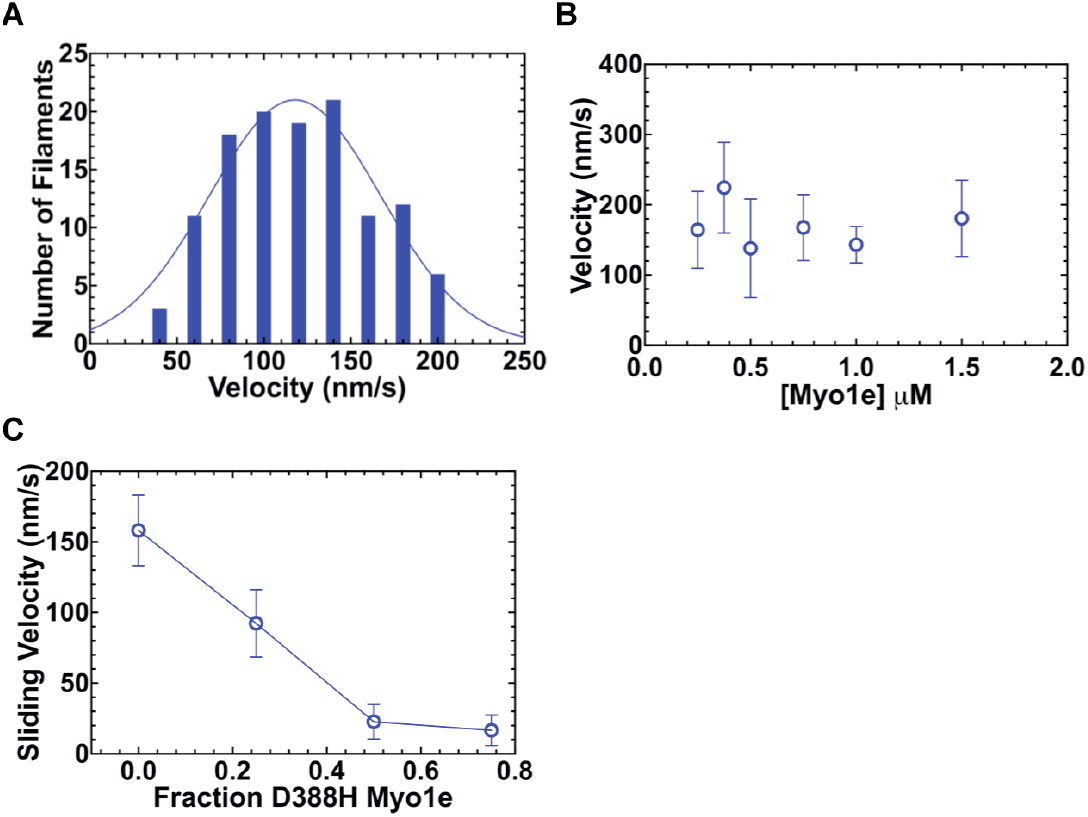
Myo1e^D388H^ lacks motor activity. **(A)** Actin filament translocation in *in vitro* motility assays using FLAG- and actin cosedimentation-purified Myo1e constructs (1μM surface density). The average sliding velocity (actin translocation speed) of Myo1e was determined by pooling the filaments from 4 independent experiments each with a separate protein preparation (*N*=120 filaments, 30 from each prep) and fitting the data to a Gaussian function. **(B)** The sliding velocity of Myo1e^WT^ was examined at a series of loading concentrations. The data is the summary of 2 independent experiments from 2 separate protein preparations. **(C)** Myo1e^WT^ was slowed by the presence of Myo1e^D388H^ in a dose-dependent manner in the mixed motility assay. The average sliding velocity is plot as a function of the fraction of Myo1e^D388H^ present ⌊(D388E/(WT + D388E)⌋ collected from 1 independent experiment.

**Table 3.** In vitro motility of Myo1e motor-IQ constructs.

| Construct | $V_{avg}$ (nm<sup>-1</sup>s<sup>-1</sup>) | # of Filaments |
| --- | --- | --- |
| WT (N=4) | 117.9±49.4 | 121 |
| D388H (N=4) | No movement | NA |

## Discussion

Association of *MYO1E* mutations with the autosomal recessive nephrotic syndrome was first identified via wholegenome linkage analysis of familial cases of SRNS [7, 51] and animal studies using Myo1e-KO mice [14, 16]. With the identification of multiple genes associated with the monogenic forms of SRNS, exome sequencing and analysis of candidate genes became a widely-used strategy for genetic diagnosis of SRNS. The assignment of new variants as pathogenic or benign can be challenging, especially in the absence of family history and detailed linkage data, and multiple lines of evidence have to be considered in deciding whether to describe a variant as potentially disease-causative [52]. Analysis of the new variants relies in part on the evolutionary conservation of the affected amino acid residues and on the variant allele frequency in large control populations. However, it can still be challenging to unequivocally evaluate the pathogenicity of the new variants to facilitate the diagnosis and targeted treatment of primary SRNS. The optimal approach to determine if a novel gene variant is likely pathogenic would rely not only on the evolutionary and statistical considerations but also on the direct functional testing of the effects of the mutations. However, animal testing (for example, using knock-in mice to directly test the effects of mutations on renal filtration) is expensive and time-consuming while testing in cell culture can be challenging if the precise functional readout for a particular protein is not well defined. In this study we used a combination of cell biology and biochemistry approaches to establish a workflow to evaluate the deleterious effects of SRNS-associated *MYO1E* mutations. In addition to identifying sensitive readouts for Myo1e activity, we have also revealed the effects of the loss of Myo1e activity on specific podocyte functions, such as endocytosis. Thus, this work not only provides an experimental workflow that can be used for future analysis of new *MYO1E* variants but also sheds new light on the functional importance of Myo1e in podocytes.

We first examined Myo1e protein stability, based on the assumption that mutations could affect Myo1e folding and degradation. Unexpectedly, we found that the stability of the Myo1e variants varied depending on the cell type used for their exogenous expression. For example, the amount of the fulllength Myo1e^T119I^ was drastically decreased in cells containing endogenous Myo1e (WT podocytes and HEK-293 cells) compared to the Myo1e^WT^ and Myo1e^D388H^, even though this variant was expressed in the Myo1e-KO podocytes at the level comparable to the Myo1e^WT^ and Myo1e^D388H^. Similarly, our attempts to express the Myo1e^T119I^-containing Motor-IQ construct in the baculovirus system were not successful due to protein degradation. Intriguingly, when we had previously introduced a Myo1e^T119I^ equivalent mutation into the genomic copy of the fission yeast *S. pombe* class I myosin, Myo1, the mutant protein was stably expressed and did not exhibit any evidence of degradation or misfolding [53]. It is also possible that the increased mRNA expression or stability may maintain the mutant Myo1e availability in the Myo1e-KO podocytes as we detected a significant increase of EGFP-myo1e mRNA levels in the cells expressing mutant versions of Myo1e compared to the wild-type version. In terms of the susceptibility of Myo1e^D388H^ to proteolysis in the baculovirus system, the proteolytic bands remained even after the motor domain preparations were purified by cosedimentation with actin filaments and ATP elution, suggesting that this protein cleavage did not affect the ability of the Myo1e motor domain to bind actin filaments or release F-actin upon ATP binding. Thus, the proteolytic cleavage may occur on a surface loop instead of the key structural elements of the motor domain. Our findings indicate that Myo1e^T119I^ and even Myo1e^D388H^ may exhibit decreased stability in some cell types; whether this loss of stability may be sufficient to completely disrupt Myo1e activity and lead to SRNS is unclear since the mutant proteins are still expressed in podocytes.

Junctional localization analysis revealed striking differences between the Myo1e^T119I^, and Myo1e^D388H^ variants. Myo1e^T119I^ was deficient in the ability to localize to podocyte cellcell contacts. Although Myo1e^D388H^ was still enriched in the junctions, we found that Myo1e^D388H^ had lower junctional dissociation rate than Myo1e^WT^ as measured by FRAP. Podocyte slit diaphragm complexes are critically important to glomerular integrity [54, 55], and the inability of Myo1e to serve as a membrane-actin linker at the cell-cell junctions may result in the disruption of the protein scaffolding at the junctions, while decreased Myo1e junctional dissociation rate may inhibit dynamic reorganization of the junctions. Combining junctional enrichment measurements with FRAP measurements of the Myo1e dissociation rate appears to be a straightforward and reliable approach for the initial evaluation of the Myo1e mutants in podocytes.

Analysis of clathrin-dependent endocytosis provides another sensitive and quantitative readout for the functionality of the Myo1e variants. The involvement of class I myosins, such as Myo1e in mammalian cells and its yeast homologs, Myo3/5 (budding yeast) and Myo1 (fission yeast), in endocytosis is highly conserved [56]. In our previous studies, overexpression of the Myo1e-tail construct (lacking the motor domain) inhibited transferrin uptake in HeLa cells [18], and introduction of motor domain mutations into the single genomic copy of Myo1 in fission yeast inhibited endocytic vesicle internalization [53]. While internalization of endocytic cargo, such as transferrin, has been traditionally used to measure the efficiency of endocytosis, either bulk cargo internalization measurements or measurements of CCV movement away from the plasma membrane may not accurately reflect changes in the early steps of clathrin coat assembly, such as initiation or stabilization of clathrin coats [57, 58]. In the present study, detailed tracking of CCV intensities and lifetimes and Myo1e localization to CCVs enabled identification of the subtle differences between the WT and mutant Myo1e. Specifically, the extent of co-localization between Myo1e and CCVs, the duration of Myo1e association with CCVs, the density of CCVs at the plasma membrane, and the distribution of CCV lifetimes were all affected by the expression of the Myo1e variants.

Our analysis of the CCV dynamics in the Myo1e-KO cells expressing various Myo1e constructs revealed differences between the two motor domain mutants as well as some unexpected contributions of Myo1e to clathrin-dependent endocytosis. Previously, based on the timing of Myo1e recruitment to CCVs (during the later stages of endocytic coat assembly and internalization [19]) and its interactions with dynamin and synaptojanin [18] and with actin assembly regulators [20, 59], Myo1e was thought to act solely as a component of the vesicle internalization and scission machinery, perhaps by helping to promote assembly of branched actin structures that facilitate vesicle internalization. Indeed, deletion of Myo1e homologs in budding and fission yeast slows down the endocytic pit invagination and actin assembly [60, 61]. Based on this proposed function, in cells lacking active Myo1e, we expected to observe prolonged CCV lifetimes, which typically represent less efficient scission, and the presence of a large fraction of stable (non-internalizing) CCSs. Indeed, we found that the average lifetimes of Myo1e-containing CCVs were longer in the cells expressing Myo1e^T119I^, and the fraction of stable CCVs (lifetime longer than 5 min) among all Myo1e-containing CCVs was increased in the Myo1e^T119I^-expressing cells. These findings are consistent with the observation that Myo1e^T119I^ exhibits weak interactions with the CCVs and, therefore, may not be able to contribute to scission. However, we also made some unexpected findings: the absence of Myo1e or the expression of Myo1e^T119I^ were associated with the lower density of CCVs on the plasma membrane. These observations suggest that Myo1e may contribute to the initial steps in clathrin coat assembly or stabilization so that in the absence of Myo1e, clathrin coat assembly is impaired. Indeed, a recent survey of the functional effects of the knockdowns of endocytic accessory proteins that relied on the automated CCV analysis placed Myo1e into the group of accessory proteins that regulate CCV stabilization [57], suggesting that sensitive approaches to the analysis of CCV dynamics can reveal novel roles for some endocytic proteins, such as Myo1e. Unlike the Myo1e^T119I^, Myo1e^D388H^ expression was associated with the increased CCV density and intensity, indicating that prolonged interactions of Myo1e with the CCVs may result in an increase in CCV initiation rate, clathrin coat stability, or the depth of endocytic invaginations **(Fig. 7)**. Thus, overall, decreased interactions between Myo1e and CCVs result in a decrease in both CCV assembly (reduced density) and internalization (extended lifetimes) while prolonged interactions lead to the increased density and intensity of the CCVs.

**Figure 7.**
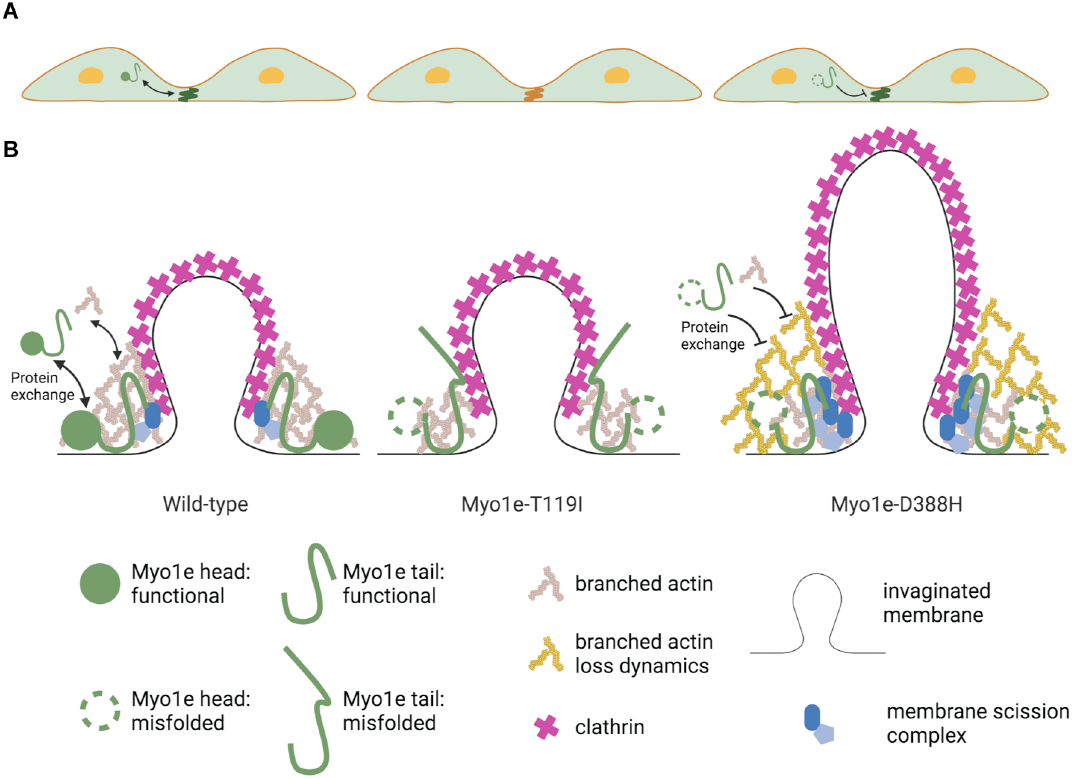
Model describing how expression of Myo1e variants affects cell junctions and clathrin-dependent endocytosis. Myo1e^WT^ transiently localizes to the boundary between plasma membrane and branched actin at the **(A)** cell-cell junctions and **(B)** clathrin-coated pits where it interacts with its binding partners (ZO-1, dynamin, synaptojanin). Myo1e tail domain interactions may promote clustering or activation of its binding partners, including phospholipids and endocytic accessory proteins. Myo1e^T119I^ is a misfolded/inactive mutant, which is rarely recruited to the CCVs or cell contacts. In the absence of Myo1e recruitment, the lifetimes of the CCVs are extended, and the number of CCVs is reduced, indicating that Myo1e^T119I^ fails to support vesicle maturation, invagination, and scission. Myo1e^D388H^ is a motor activity-deficient mutant characterized by prolonged retention at the cell-cell junctions and CCVs. Since its tail domain is still functional and capable of interacting with its binding partners, expression of this mutant results in increased accumulation of endocytic accessory proteins, leading to higher CCVs density and intensity than expression of the Myo1e^WT^.

ATPase activity measurements and *in vitro* motor activity assays represent the most direct tests of the effects of mutations on the motor domain functions. Previously, Myo1e ATPase activity was characterized using a purified protein consisting of the motor and neck domains [50] while purification of the full-length Myo1e has been challenging. The ability of Myo1e to translocate actin filaments has never been measured. We were able to measure Myo1e-driven actin motility for the first time, resulting in an estimate of the actin sliding speed of ~120 nm/sec, which is faster than some of the short-tailed class I myosins, Myo1a, with the actin sliding speed of 28 nm/sec [62] and Myo1b, with the speed of 56 nm/sec [63] and comparable to the short-tailed Myo1c at 83 nm/sec [64]. As expected, Myo1e exhibited actin-activated ATPase activity. The high basal ATPase activity and maximal ATPase activity of Myo1e^WT^ is consistent with the previous kinetics study [50], but the concentration of actin required to activate ATPase activity is 10-times higher in this study than that of the previous study (*K*_ATPase_=10.7±7.5 μM in this study vs. 1.2±0.36 μM in [50]). The buffer in the current study contained a 2-fold higher MgCl_2_, which can have an impact on myosin ATPase kinetics [33]. The dramatic reduction in both basal and actin-activated ATPase activity of Myo1e^D388H^ indicates a severe defect in the ATPase cycle of this mutant. The results of single ATP turnover measurement are consistent with the steady-state ATPase assay for the Myo1e^WT^ and Myo1e^D388H^, supporting the conclusion of the deficient but not completely abolished ATPase activity of Myo1e^D388H^. Previously, it has been shown that the D454A mutation in the *Dictyostelium* myosin II (D454 being an equivalent residue to D388 in Myo1e) results in the loss of motor activity in the sliding filament assay, indicating that this aspartate residue in the switch-2 region is essential for myosin motor activity in both class I and class II myosins [65].

Overall, our analysis identified differences between the two mutants tested and between the mutants and the wild-type Myo1e **(Fig. 7)**. While Myo1e^T119I^ rarely co-localized with CCVs or cell-cell junctions and maintained its association with the CCVs for only short time periods, Myo1e^D388H^ remained associated with the CCVs and cell junctions longer than Myo1e^WT^. What could account for these differences in intracellular localization and dynamics? One explanation is that actin binding via the motor domain or actin-dependent transport are required for Myo1e recruitment to its sites of action, and that Myo1e^T119I^ lacks the ability to bind actin, which we were not able to test directly due to its proteolysis during purification. However, our previous studies show that a Myo1e tail construct (lacking the motor and neck domains) is enriched in the cell-cell junctions and CCVs [17, 18], arguing against the direct contribution of the motor domain to localization. Second, the T119I mutation could lead to complete misfolding, including misfolding of the tail domain, preventing the tail from performing its normal function in Myo1e localization. Third, while it is not known whether Myo1e motor activity is regulated via intramolecular interactions, other myosins have been shown to be autoinhibited via tail-motor binding [66]. If this mode of regulation is also applicable to Myo1e, the mutant motor domain may not be able to undergo the conformational change required to relieve the auto-inhibition. On the other hand, Myo1e^D388H^ exhibited prolonged association with cell junctions and CCVs. One potential explanation is that this mutation reduces the rate of ADP dissociation or slows dissociation from actin in the presence of ATP, which could prolong actin binding. However, both Myo1e^WT^ and Myo1e^D388H^ were able to dissociate from actin in the presence of ATP and have a similar *K*ATPase, suggesting Myo1e^D388H^ is not a true rigor mutant. Alternatively, if the auto-inhibition model for regulation of Myo1e activity is correct, perhaps the D388H mutation allows Myo1e to remain in an open (activated) conformation longer, thereby prolonging tail domain interactions with its binding partners. Our finding that expression of the D388H variant increases CCV density at the plasma membrane suggests that prolonged tail domain interactions with the plasma membrane phospholipids or endocytic accessory proteins promote assembly and maturation of CCVs, suggesting a new role for the myosin tail domain.

In summary, in this study we have developed a pipeline for characterization of the Myo1e mutations, which can be helpful in genetic diagnosis of SRNS, prioritizing the clinical decisions, and, potentially, in the development of Myo1e-targeted treatment. We have also identified Myo1e^T119I^ and Myo1e^D388H^ as likely pathogenic variants associated with SRNS.

## End Matter

### Author contributions

P.L., L.G., C.Y., D.P., J.B.K., C.P., S.C., M.P., E.P. and C.M. performed experiments; P.L. and L.G. analyzed data; P.J., C.Y., F.H., and M.K. edited and revised manuscript; P.L., C.Y., and M.K. interpreted results of experiments; P.L., S.L., C.Y., F.H., and M.K. were involved in conception and design of research; P.J., L.G. and C.Y. prepared figures; P.J. and M.K. drafted manuscript; P.J. and M.K. approved final version of manuscript.

## Acknowledgments

We are grateful for the critical discussions with David W Pruyne, Peter Calvert and Mehdi Mollapour throughout the project development. We thank Sarah Barger, Michael Garone, and Jacqualyn Schulman for the early discussion of the project and assistance with cell culture, Western blotting and imaging analysis, Matthew J. Gastinger (Oxford Instruments, Inc.) for help using Imaris, Wayne A. Decatur and Sean M. Lantry (Osmose Utilities Services, Inc.) for generously creating data analysis tools. Illustrations were created using BioRender.com.

---

**Supplementary Figure 1.**
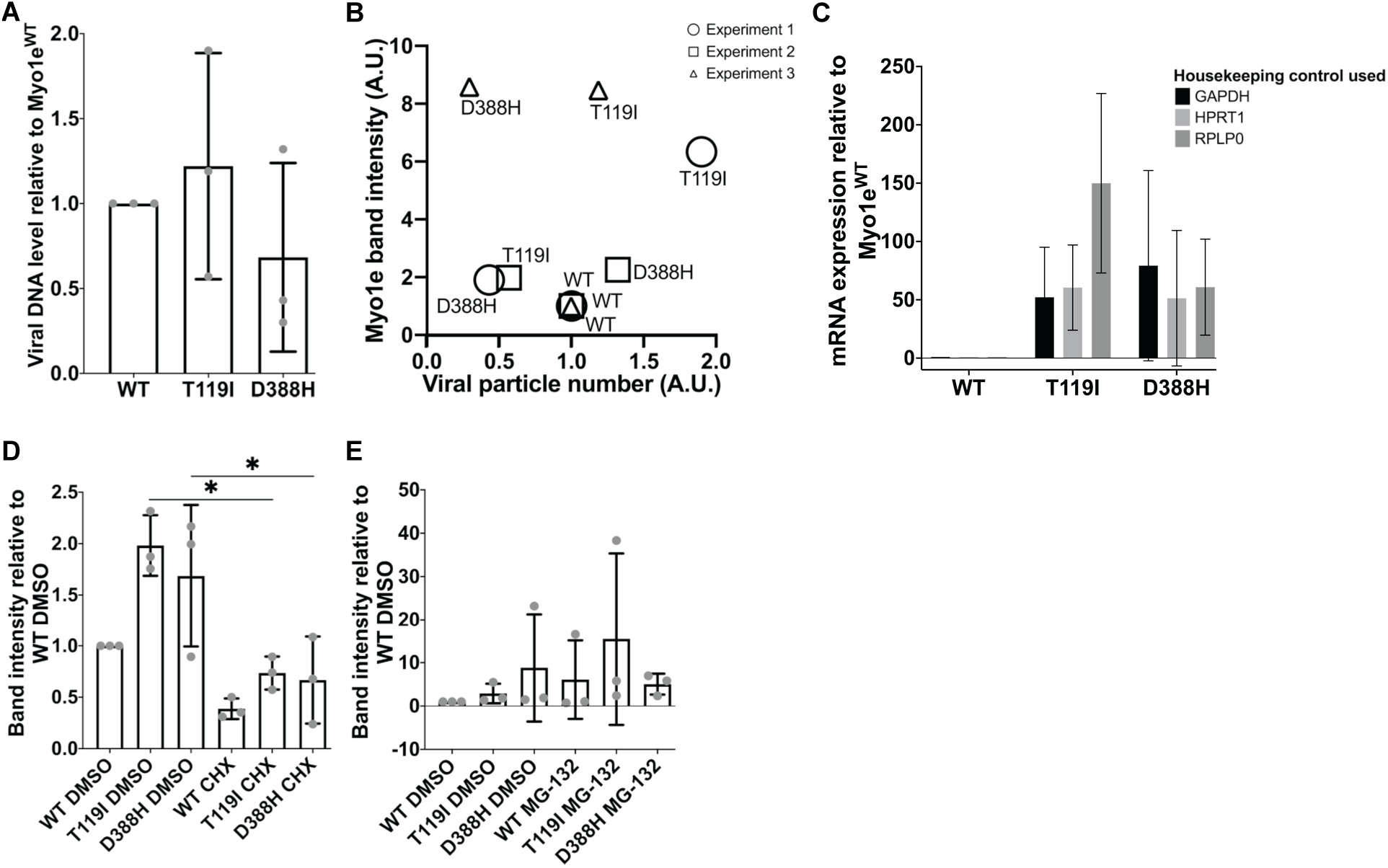
Expression of recombinant Myo1e constructs in Myo1e-KO podocytes using adenoviral vectors (related to Figure 2). **(A)** The efficiency of the adenoviral delivery of the various EGFP-Myo1e constructs into podocytes was compared by performing quantitative PCR-amplification of the EGFP coding sequence using viral DNA isolated from infected cells as a template. The values obtained for EGFP was normalized to genomic GAPDH and mutant values to those for the WT. **(B)** Analysis of correlation between viral DNA amount (determined as in A) and Myo1e protein level (determined from Western blot band intensity) indicates that the higher viral load in the cell does not directly correlate with the higher protein band intensity in these experiments. **(C)** EGFP-Myo1e construct mRNA expression was quantified by quantitative RT-PCR. EGFP-Myo1e RNA amount was determined by normalization of the values for EGFP to those for three housekeeping genes, GAPDH, HPRT1 or RPLP0, and the resulting number for the mutants was normalized to the WT in each experiment. **(D)** Related to Fig. 2C-D. Myo1e protein level after CHX treatment of the Myo1e-KO podocytes expressing various EGFP-Myo1e constructs. EGFP-Myo1e band intensity was normalized to that for the EGFP-Myo1e^WT^ in DMSO-treated cells. **(E)** Related to Fig. 2E-F. Myo1e accumulation level after MG-132 treatment of the Myo1e-KO podocytes expressing various EGFP-Myo1e constructs. Band intensity was normalized to that for the EGFP-Myo1e^WT^ in DMSO-treated cells. **(A-E)** Data collected from 3 independent experiments. Data shown in A, C, D, E are mean±SD. Asterisks indicate a significant difference from Myo1e^WT^ or indicated controls as determined by one-way ANOVA, *P≤0.05.

**Supplementary Figure 2.**
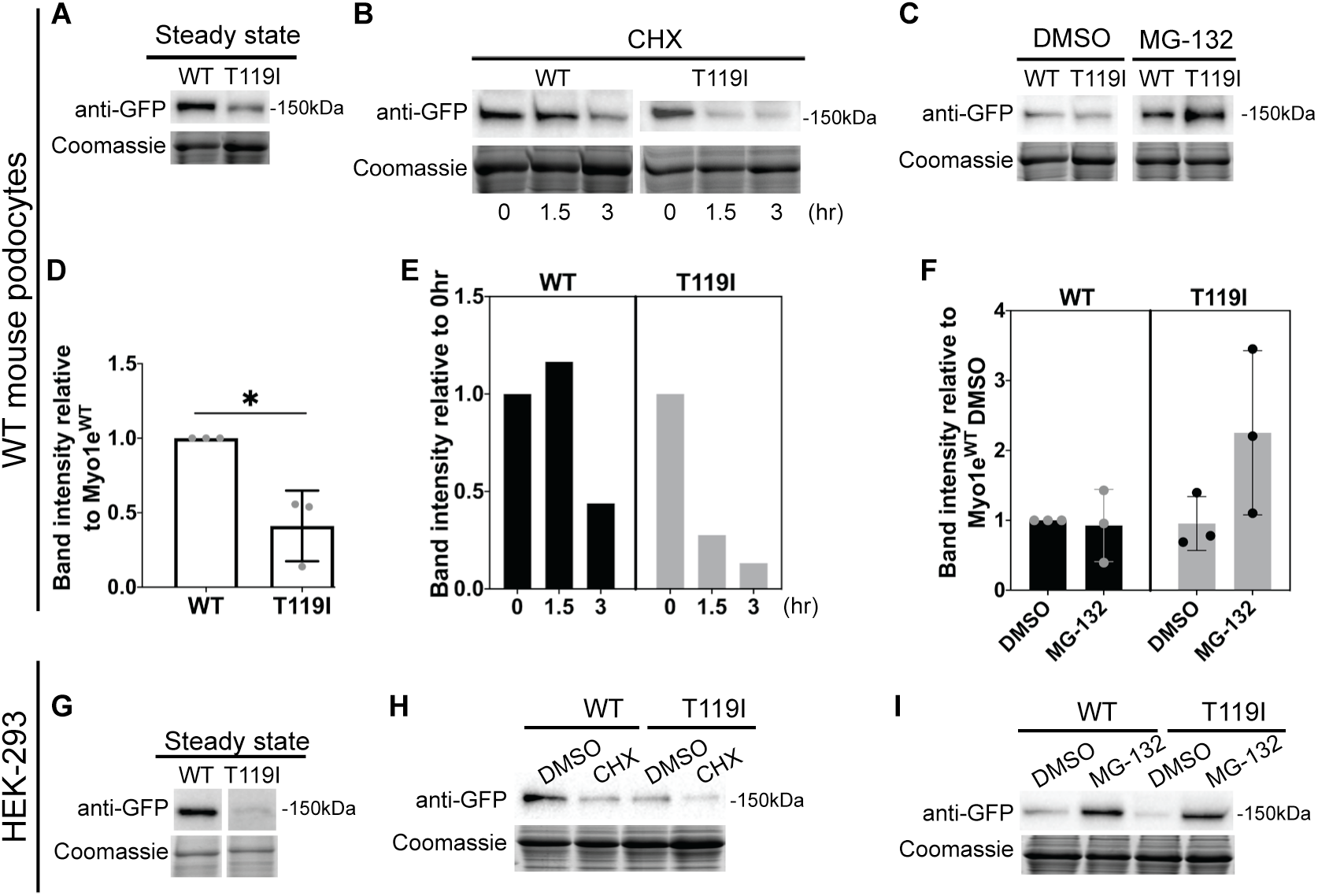
Myo1e^T119I^ is unstable in the cell lines with endogenous Myo1e. **(A)** Western blot analysis of total cell lysates of WT podocytes expressing Myo1e^WT^ and Myo1e^T119I^ at steady state. **(B)** Western blot analysis of total cell lysates of WT podocytes expressing Myo1e^WT^ and Myo1e^T119I^ following 20 μg/ml cycloheximide CHX) treatment. **(C)** Western blot analysis of total cell lysates of WT podocytes expressing Myo1e^WT^ and Myo1e^T119I^ following 10 μM MG-132 treatment for 3-4 hours. **(D)** EGFP-Myo1e band intensity normalized to that for the WT (as in A) (mean±SD). Data collected from 3 independent experiments. There is a significant decrease in the intensity of the Myo1e^T119I^ band compared to the Myo1e^WT^ as determined by one-way ANOVA, *P≤0.05. **(E)** Quantification of the EGFP-Myo1e band intensity following CHX treatment (as in B). The intensity for each construct was normalized to the intensity for the same construct at time 0. Data collected from 1 experiment. **(F)** Quantification of the EGFP-Myo1e band intensity after MG-132 treatment (as in C) and normalized to the Myo1e^WT^ DMSO (mean±SD). Data collected from 3 independent experiments with 3 (2 repeats) or 4 hours (1 repeats) of MG-132 treatment. **(G)** Western blot analysis of total cell lysates of HEK-293 cells expressing Myo1e^WT^ and Myo1e^T119I^ at steady state. **(H)** Western blot analysis of total cell lysates of HEK-293 cells expressing Myo1e^WT^ and Myo1e^T119I^ following 0.5 μg/ml cycloheximide (CHX) treatment for 6 hours. **(I)** Western blot analysis of total cell lysates of HEK-293 cells expressing Myo1e^WT^ and Myo1e^T119I^ following 10 μM MG-132 treatment for 24 hours. **(A-C, G-I)** Blots were probed with anti-GFP antibody. Equal protein loading verified by Coomassie Blue staining.

**Supplementary Figure 3.**
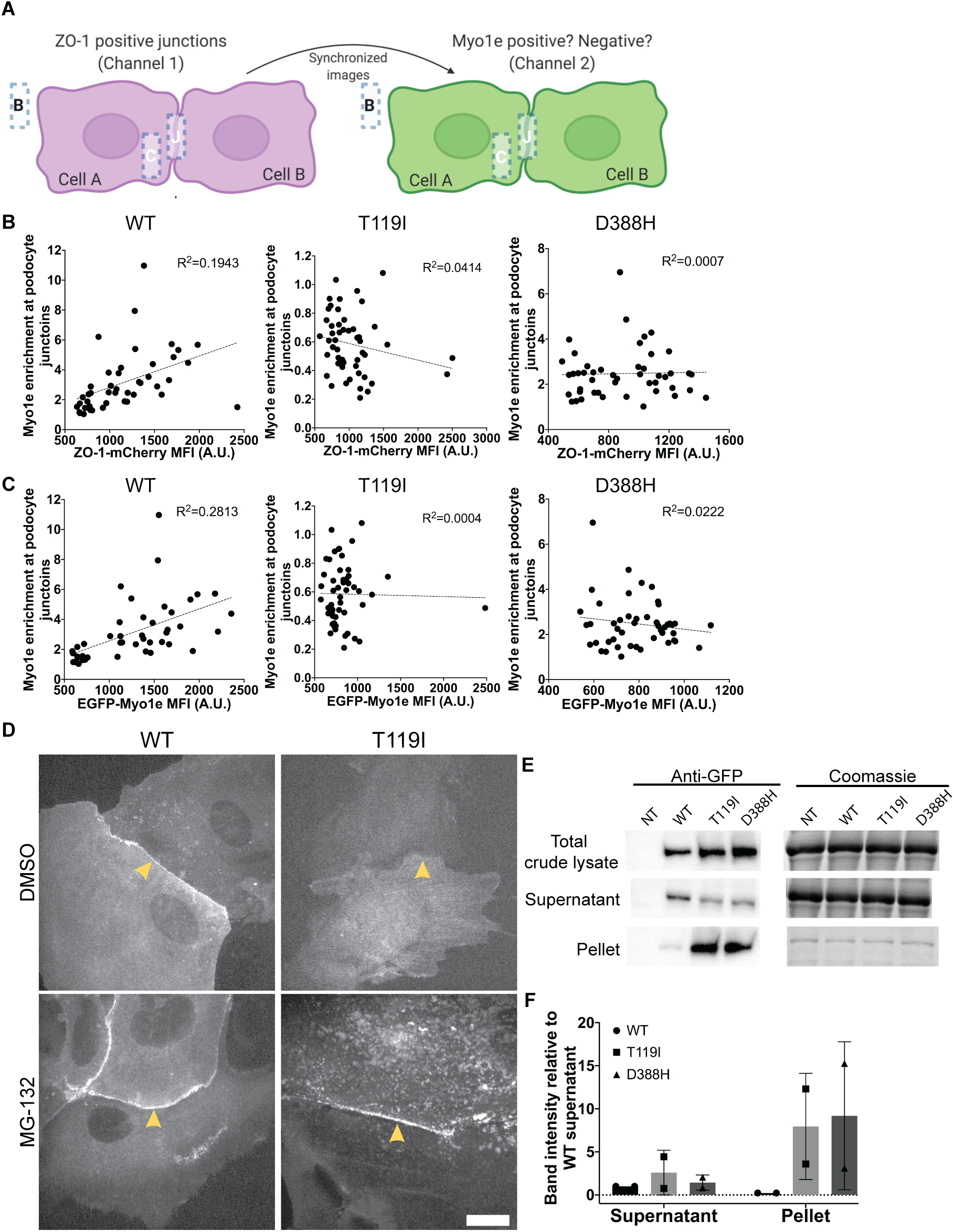
Analysis of EGFP-Myo1e localization at cell-cell junctions in the Myo1e-KO podocytes (related to Figure 3). **(A)** An illustration of the junctional enrichment analysis method for live-cell images in Fig. 3A-B. Junctional enrichment was determined by dividing EGFP fluorescence intensity at the cellcell contacts (J) by the cytosolic EGFP intensity (C) after subtracting the non-cell (background) intensity (B). To be unbiased in the junctional ROI selection, we used ZO-1-mCherry channel images, rather than EGFP-Myo1e images, to identify the junctions. **(B-C)** Correlation plots of Myo1e enrichment at podocyte junctions vs. mean fluorescence intensity (MFI) of ZO-1-mCherry (**B)** or EGFP-Myo1e (**C**). The dotted lines indicate linear regression fit. The linear regression R square values are shown in the graphs. Each data point indicates a single cell. **(D)** A single confocal section of Myo1e-KO podocytes expressing EGFP-Myo1e^WT^ or -Myo1e^T119I^ with DMSO or 10 μM MG-132 treatment for 3 hours. Yellow arrowheads indicate cell-cell junctions. Scale bar, 20 μm. **(E)** Western blot analysis of total cell lysates, supernatants and pellets of KO podocytes non-infected (NT) or expressing Myo1e^WT^, Myo1e^T119I^ and Myo1e^D388H^ following 10 μM MG-132 treatment for 3 hours. Blots were probed with anti-GFP antibody. Equal protein loading verified by Coomassie Blue staining. **(F)** Quantification of band intensity in (E) normalized to the Myo1e^WT^ supernatant (mean±SD). Data collected from 2 independent experiments.

**Supplementary Figure 4.**
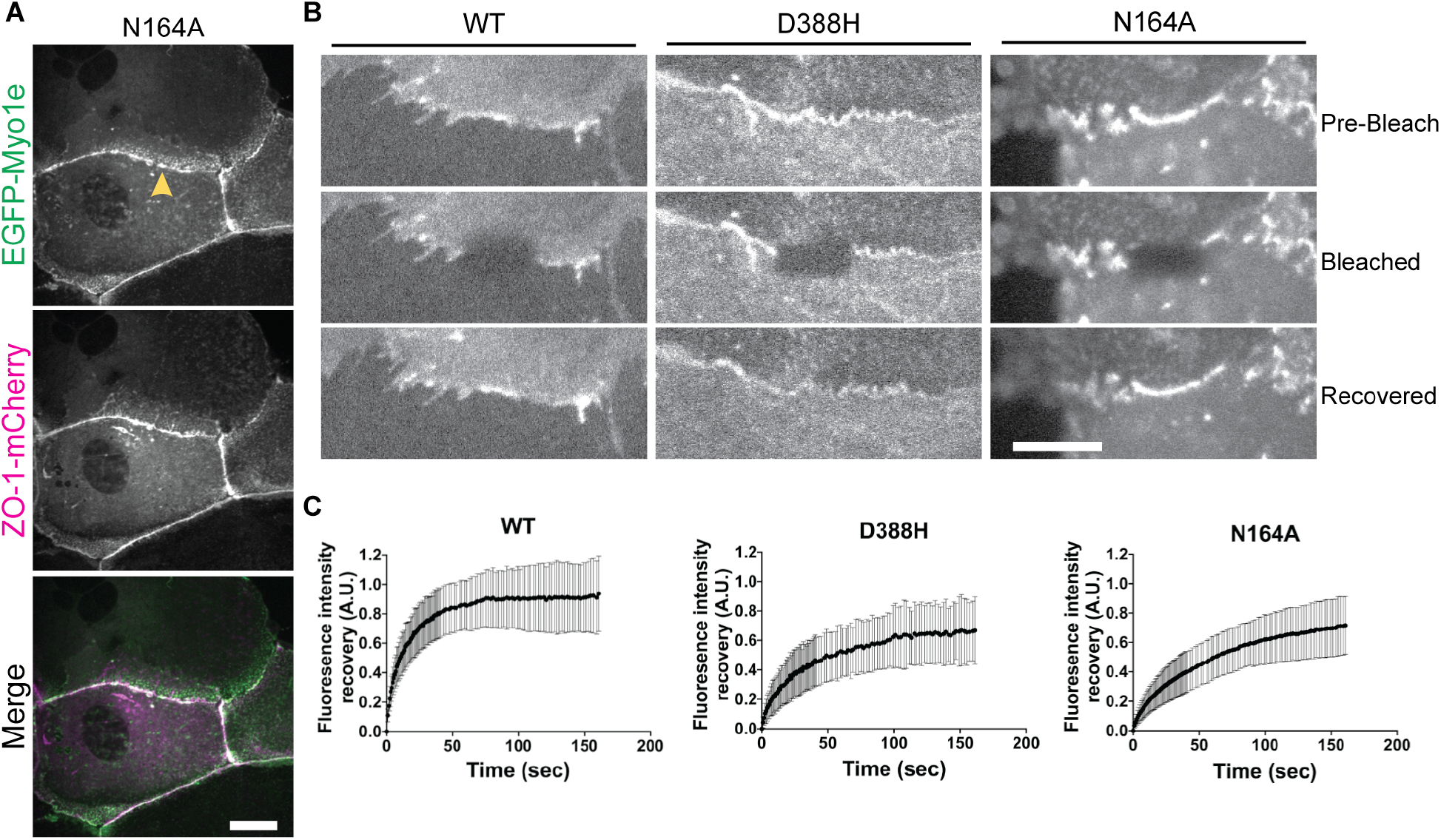
Myo1e^D388H^ exhibits slower protein exchange at the podocyte junctions (related to Figure 3). **(A)** A single confocal section of Myo1e-KO podocytes co-expressing ZO-1-mCherry and EGFP-Myo1e^N164A^. Yellow arrowhead indicates cell-cell junction. Scale bar, 20 μm. **(B)** Three time frames (single confocal sections) from the time-lapse movies of FRAP analysis of the Myo1e-KO podocytes expressing Myo1e^WT^, Myo1e^D388H^, or Myo1e^N164A^. Scale bar, 10 μm. **(C)** FRAP curves (mean±SD) of 26 Myo1e^WT^-, 33 Myo1e^D388H^- and 33 Myo1e^N164A^-expressing cells from 2 to 3 independent experiments. The intensity data were corrected for photobleaching and normalized to the pre-bleach intensities.

**Supplementary Figure 5.**
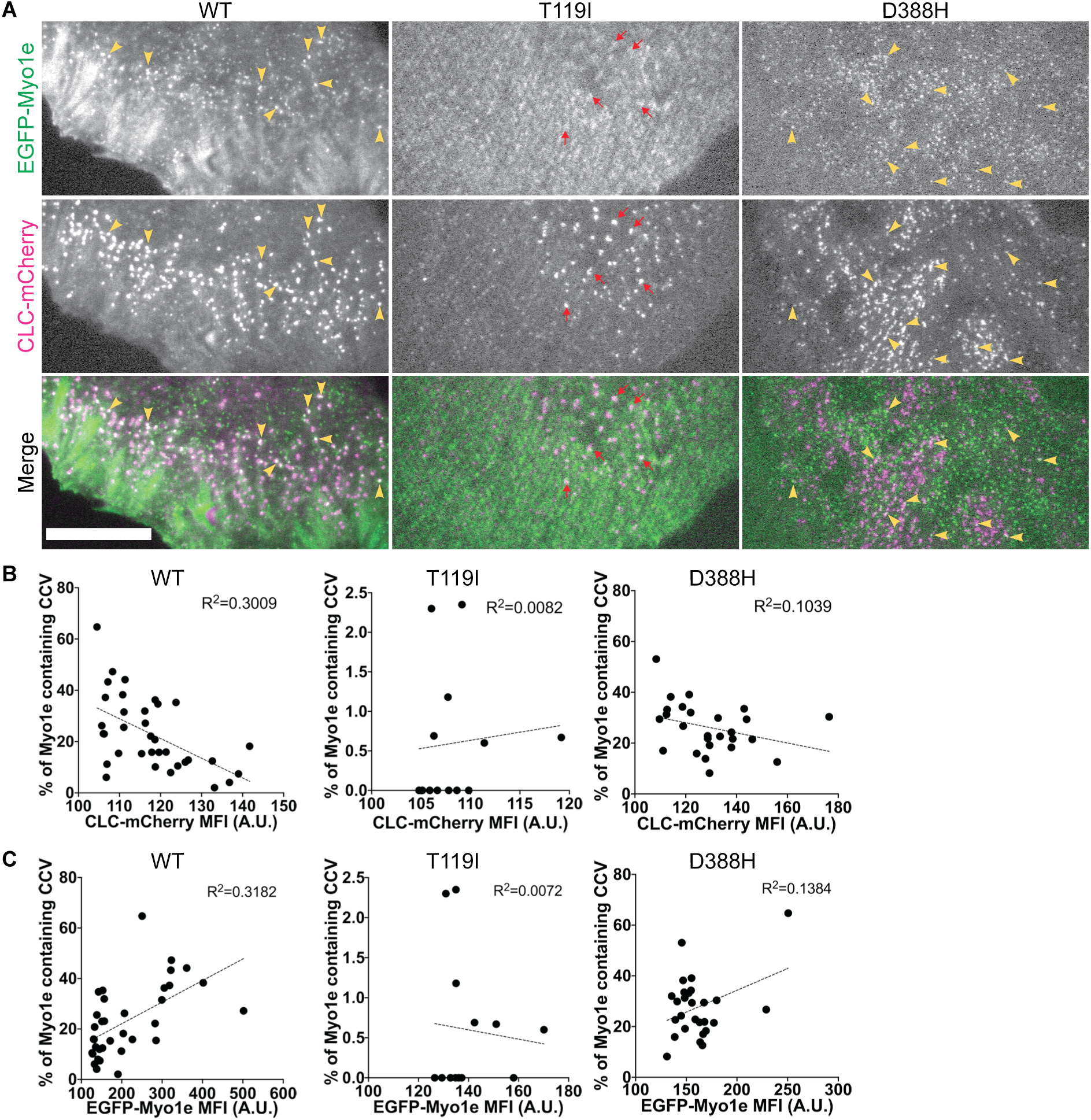
Analysis of EGFP-Myo1e co-localization with CCVs by TIRF microscopy (related to Figure 4). **(A)** TIRF images of Myo1e-KO podocytes co-expressing clathrin-light-chain-mCherry (CLC-mCherry) and EGFP-Myo1e^WT^, -Myo1e^T119I^ or -Myo1e^D388H^. Yellow arrowheads point to Myo1e-containing CCVs while red arrows mark CCVs without Myo1e co-localization. Scale bar, 20 μm. **(B-C)** The correlation plots of percent of CCV-Myo1e co-localization vs. mean fluorescence intensity (MFI) of CLC-mCherry (**B**) or of EGFP-Myo1e (**C**), along with the linear regression lines (dotted line). The linear regression R square values are shown. Each data point indicates a single cell.

**Supplementary Figure 6.**
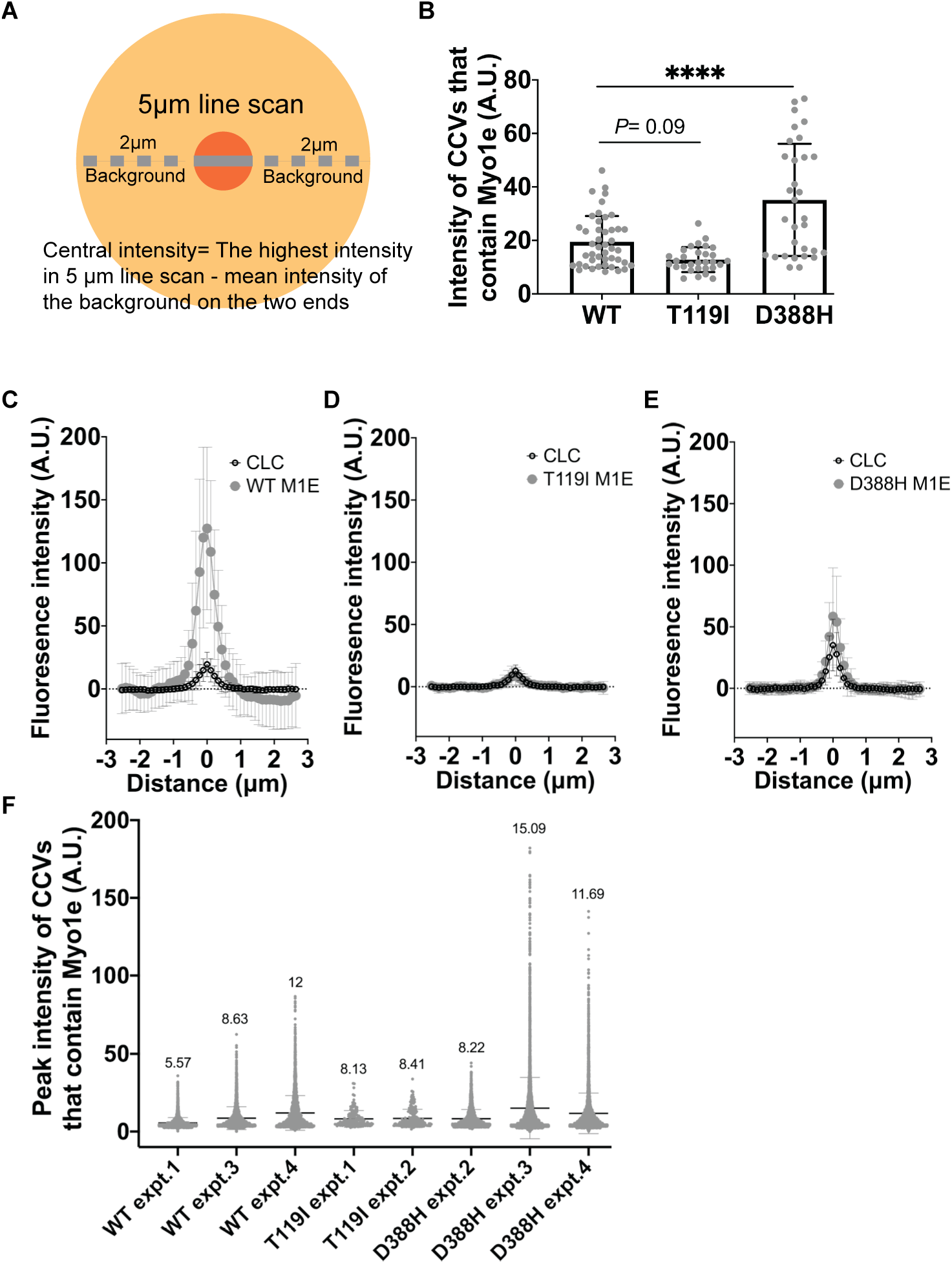
Analysis of the fluorescence intensity of CCVs co-localized with Myo1e (related to Figure 4). **(A)** An illustration of the line scan method used to measure the intensity at the center of each CCV (central intensity). The highest intensity value within the 5 um-long line scan across each CCV was corrected by the local background around the CCV. **(B)** Central fluorescence intensity (mean±SD) of the CCVs that contain Myo1e as measured by line scans and background-subtracted as shown in (A). CCVs from 9 Myo1e^WT^ (N=45), 6 Myo1e^T119I^ (N=30), and 6 Myo1e^D388H^ (N=31) expressing cells from 2 to 3 independent experiments were quantified. **(C-E)** Fluorescence intensity curves (mean±SD) derived from line scans across EGFP-Myo1e-containing CLC-mCherry CCVs. **(C)** EGFP-Myo1e^WT^ **(D)** EGFP-Myo1e^T119I^ **(E)** EGFP-Myo1e^D388H^. **(F)** Peak fluorescence intensity (mean±SD) of the CCV tracks that contain Myo1e is shown in batches corresponding to each individual experiment. Mean value of each experimental group is shown. The panel is related to Fig. 4C.

**Supplementary Figure 7.**
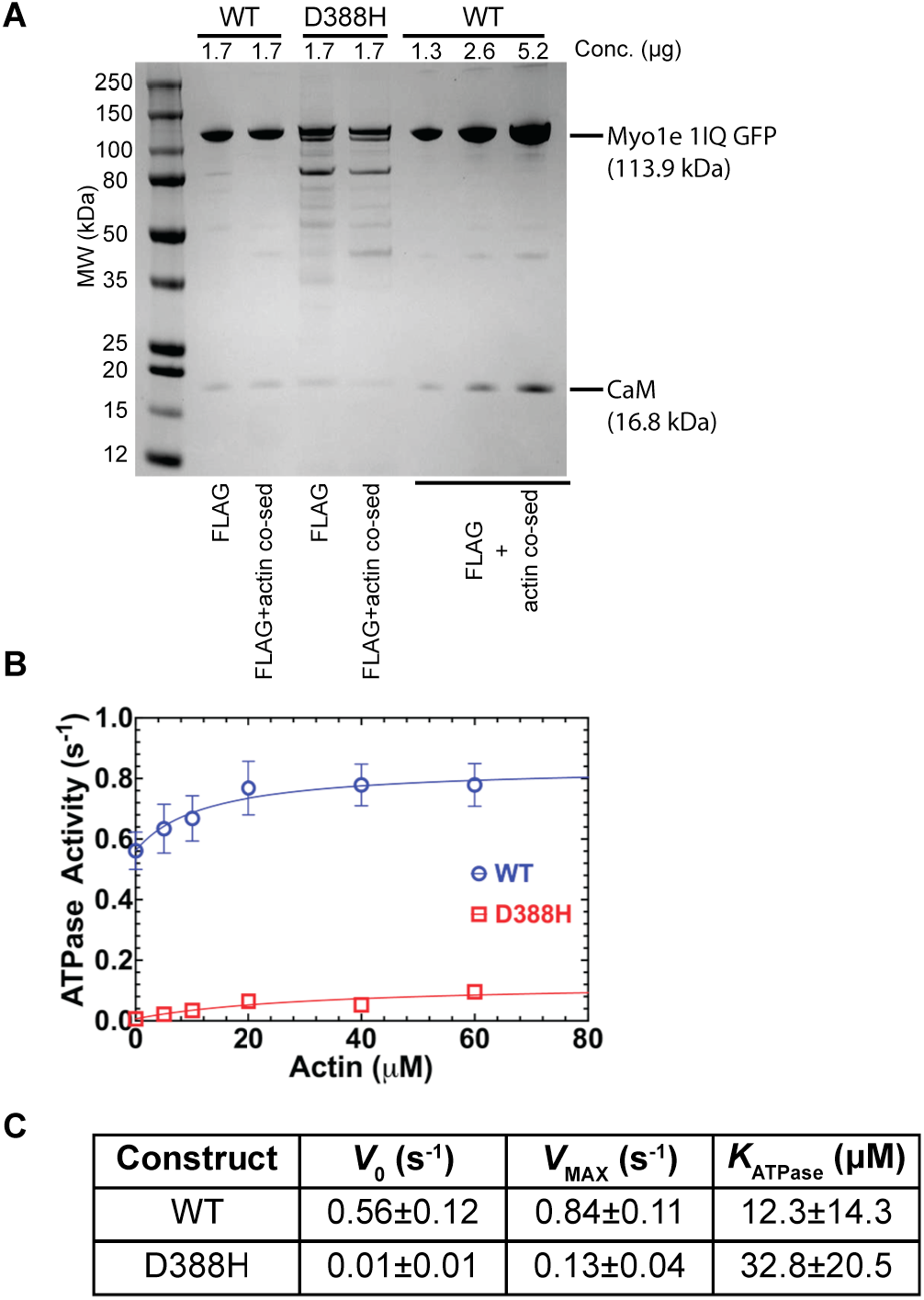
Purification of Myo1e^WT^ and Myo1e^D388H^ constructs for in vitro motor activity experiments. **(A)** Coomassie-stained SDS-PAGE showing fractions from the purification of the Motor-IQ domain constructs (Myo1e wild-type and D388H) using the baculovirus expression system. **(B)** ATPase activity measurements using FLAG-purified Myo1e constructs. **(C)** Summary of ATPase values determined with FLAG-purified Myo1e constructs.

**Supplementary table 1.**
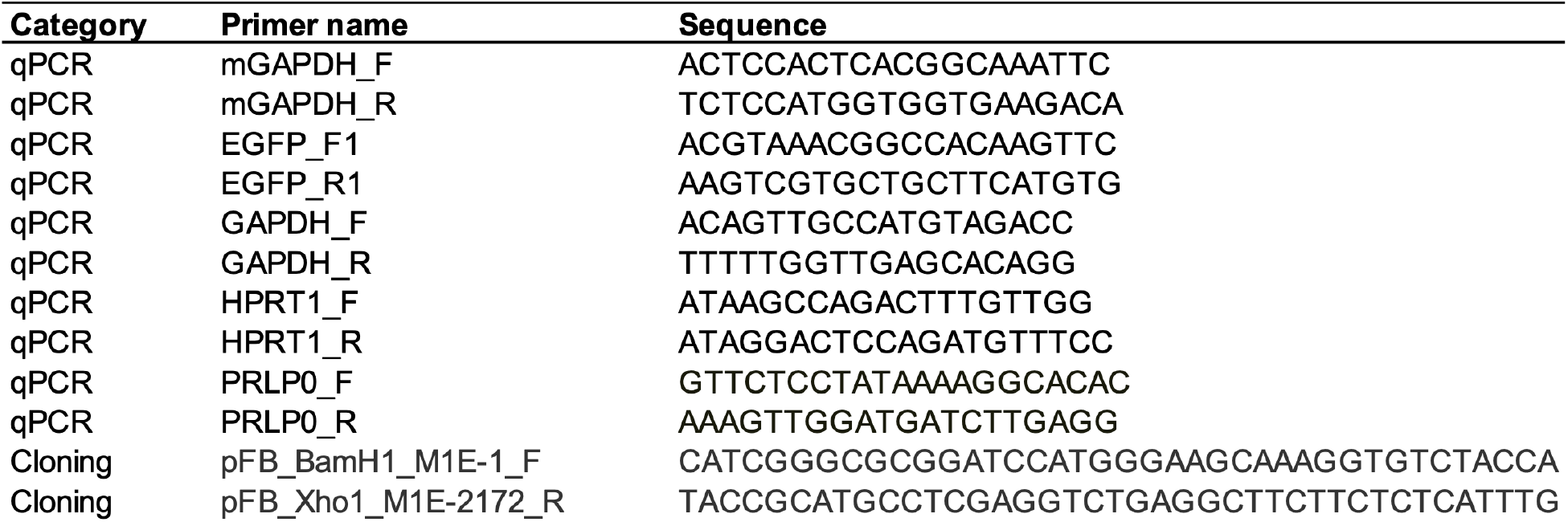
List of primer sequences used in this paper.

**Supplementary table 2.**
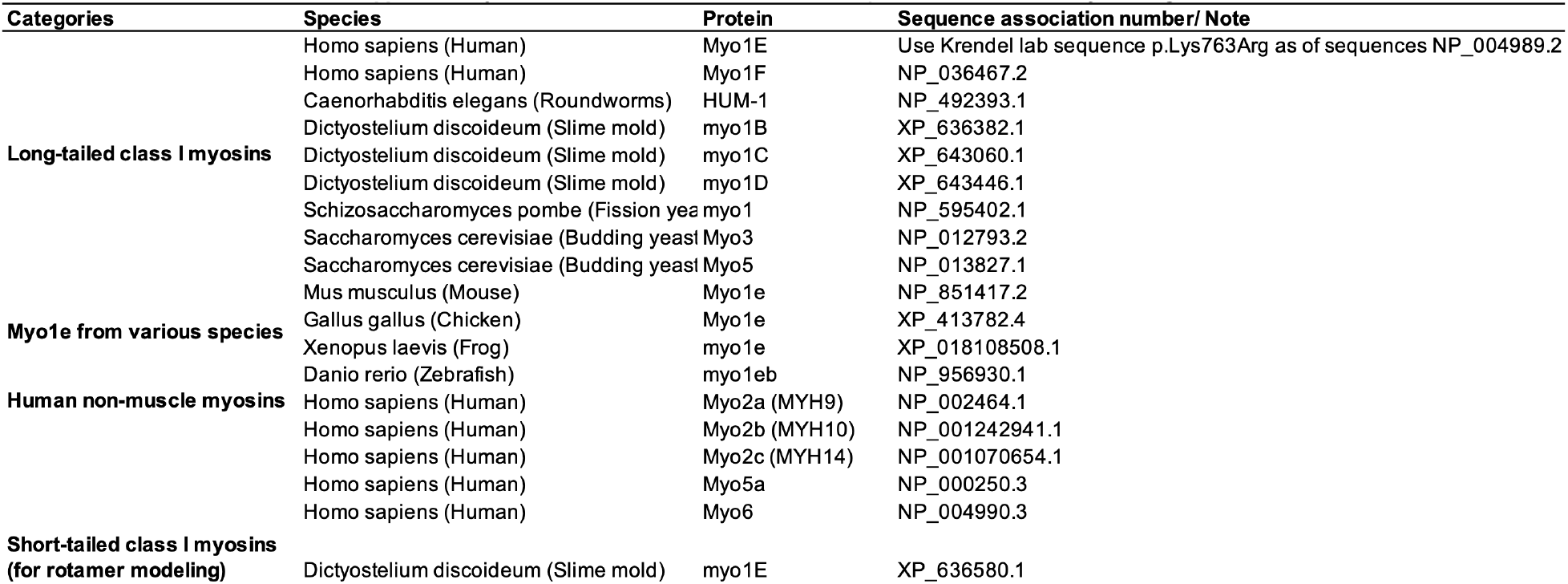
Accesion numbers for the sequences used for the Myo1e alignments.

**Supplementary table 3.**
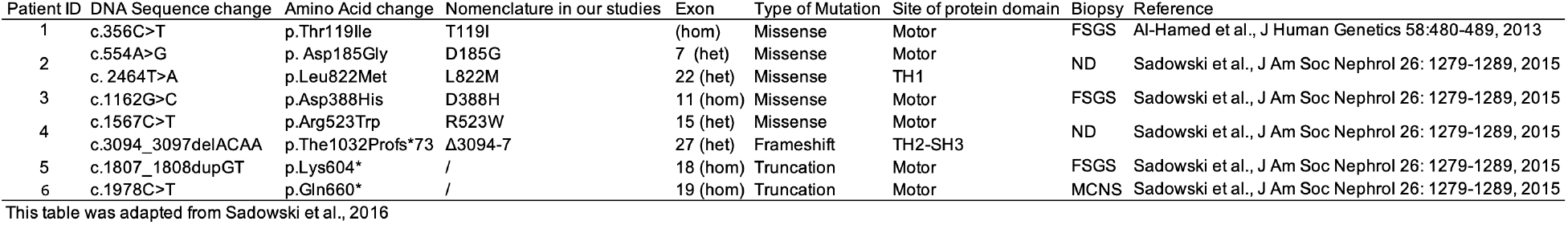
Summary of the recently identified *MYO1E* variants.

## Supplementary movies

**Supplementary movie 1 (related to figure 3E&F, S4B&C). Representative movies of fluorescence recovery after photobleaching (FRAP) at the Myo1e-KO podocyte junctions.** (**Left)** Myo1e^WT^. **(Right)** Myo1e^D388H^. 5 frames of pre-bleaching images were collected at 1 frame/sec. Post-bleaching images were acquired at the rate of 2 frames/sec for the first 40 sec and 1 frame every 2 sec for the subsequent 120 sec. The frame rate of the movie is 20 frames per second for a total of 117 frames. Scale bar, 10 μm.

**Supplementary movie 2. Representative movies of F-actin translocation by Myo1e motor+IQ (related to figure 6).** (**Left)** Myo1e^WT^ motor+IQ. **(Middle)** Myo1e^D388H^ motor+IQ. **(Right)** Myo1e^WT^+ Myo1e^D388H^ motor+IQ. The frame interval for the imaging is 10 seconds with 10 minutes duration. The frame rate of the movie is 20 frames per second for a total of 61 frames. Scale bar, 10 μm.

## Supplementary materials

**Protein expression and drug treatments (related to figure S2).** To measure protein expression, wild-type podocyte lysates were collected 48 hrs post-infection. Protein degradation rate was measured after 1.5- or 3-hour treatment with 20μg/ml of cycloheximide (Sigma-Aldrich #C7698) in the complete RPMI medium. Protein accumulation rate upon proteasome inhibition was measured after 3- or 4-hour treatment with 10μM of MG-132 (Cell Signaling Technology #2194) in the RPMI medium. For HEK-293 cells, steady state protein lysates were collected 24 hrs post-transfection. Protein degradation rate was measured following 6-hour treatment with 0.5 μg/ml cycloheximide in the complete DMEM medium. Protein accumulation rate upon proteasome inhibition was measured following 10 μM MG-132 treatment for 24 hours.

**Transfection (related to figure S2).** 1 ug of Myo1e mutant constructs on the EGFP-C1 vector were transfected to HEK-293 cells in 12-well plates by using Lipofectamine 3000 transfection reagent (Invitrogen) following the manufacturer’s instructions.

